# Snapshots of ribosome dynamics near atomic-resolution in situ: full insight into the eukaryotic elongation cycle

**DOI:** 10.1101/2023.07.12.548775

**Authors:** Jing Cheng, Chunling Wu, Junxi Li, Qi Yang, Mingjie Zhao, Xinzheng Zhang

**Author notes:** Corresponding author Correspondence to Xinzheng Zhang.

## Abstract

Many protein complexes are highly dynamic in cells, characterizing their conformational changes in cells is crucial for unraveling their functions. In this report, using cryo-electron microscopy, 451,700 ribosome particles from *Saccharomyces cerevisiae* cell lamellae were obtained to solve the 60S region to 2.9 Å resolution by in situ single particle analysis. Over 20 distinct conformations were identified by 3D classification with resolutions typically higher than 4 Å. These conformations were used to reconstruct a complete elongation cycle of eukaryotic translation with elongation factors. We found that compact eEF2 anchors to the partially rotated ribosome after subunit rolling, and hypothesize that it stabilizes the local conformation for peptidyl transfer. Moreover, an open eEF3 binding to a fully rotated ribosome was observed, whose conformational change was coupled with head swiveling and body back-rotation of the 40S subunit.

## Main

Studying conformational dynamics of protein machines by cryo-electron microscopy (cryo-EM) is an important endeavor. In the past few years, structural studies of multi-conformations have been achieved primarily by in vitro single-particle analysis (SPA), with high resolutions readily obtained. However, simulating the cellular environment in vitro is challenging, resulting in issues such as not observing all macromolecular activities^1^. In situ dynamic structural studies of proteins in cells provide a full complement of protein functional states compared with SPA, but the resolution of in situ protein structures in multiple states is usually limited.

Protein synthesis performed by ribosomes is a complex process consisting of four steps: initiation, elongation, termination and ribosome recycling. Incorporating each new amino acid into the nascent peptide chain during elongation involves multiple conformational rearrangements of the ribosome. tRNA base pairs with the mRNA codon template, and ribosomes synthesize polypeptides from amino acids carried by the ribosome-bound tRNAs. The peptidyl transferase center (PTC) is located in the large subunit (LSU) of the ribosome and catalyzes peptide bond formation and transfer of amino acids carried by tRNA to the growing peptide chain. Eukaryotic elongation factor 1 (eEF1A) combined with GTP facilitates the transport of the aminoacyl-tRNA to the A/T site, and eukaryotic elongation factor 2 (eEF2) assists in the translocation of the ribosome along the mRNA reading frame to ensure fidelity. The tRNA at the A- and P-sites also translocate to the P- and E-sites, respectively, and the vacated A-site receives the aminoacyl tRNA required for the next round of peptide chain extension. Deacyl-tRNA at the P-site is then released from the ribosome, and particular eukaryotes, such as *Saccharomyces cerevisiae* (*S. cerevisiae*)^2^, also require eukaryotic elongation factor 3 (eEF3) for this process.

During ribosomal translation, ribosomes, tRNA and mRNA undergo large-scale conformational changes. Some intermediate states have been analyzed by X-ray crystallography and cryo-EM (reviewed in ^3^), but the complexity of eukaryotic systems has limited this analysis, and therefore, many in vitro studies have focused on bacteria^4,5^. In addition, recent studies based on cryo-electron tomography have elaborated on cellular translation processes in eukaryotes^6,7^ and prokaryotes^8^. However, these studies have only achieved high-resolution structural analysis of an averaged ribosome in different states, with the resolution of individual states limited by the number of particles used, resulting in an average resolution of 6 to 7 Å, which is insufficient for atomic model building. In addition, these studies may not observe conformations with relatively small populations when analyzing protein dynamics.

In this report, to efficiently achieve a large dataset of ribosome particles inside cells, we investigated the eukaryotic ribosome of *S. cerevisiae* from cryo-focused ion beam (FIB) milled cells by using the in situ single particle analysis (isSPA) method^9,10^ and determined 24 distinct in situ ribosome structures at resolutions ranging from 3.2 to 4.8 Å. Notably, most of these structures were complexed with eukaryotic elongation factors (eEFs), with several of these states previously unresolved. We reconstructed the elongation cycle using these high-resolution structures to present a complete eukaryotic elongation cycle.

## Results

### In situ structure determination of multiple ribosome conformations

*S. cerevisiae* lamellae (Fig. 1a, b) were thinned by cryo-FIB milling to visualize various conformations of the eukaryotic ribosome during translation. With a total of 451,700 filtered ribosome particles detected by GPU accelerated isSPA (GisSPA)^11^ from ∼40 lamellae, we achieved an overall resolution of 2.9 Å for the 60S subunit (Fig. 1c). The high-resolution map revealed the well-resolved bases of ribosomal RNA (rRNA) and amino acid side chains of the ribosomal proteins (Fig. 1d). The local resolution of 80S varies noticeably between the 40S and 60S regions because of 40S subunit structural dynamics. Thus, individual conformations were identified. To identify the multiple states in translation and study structural heterogeneity, we performed focused 3D classifications based on well-refined 80S ribosome particles without alignment using masks to include regions with heterogeneities via cryoSPARC^11^ (see Methods). The 3D classification analysis yielded 24 different conformations (Fig. 2 and Extended Data Figs. 1 and 2) differentiated by the 40S subunit, tRNAs, eEF2, eEF3 and the L1 stalk. The resolution of each class is listed in Supplementary Table 1.

**Fig 1.**
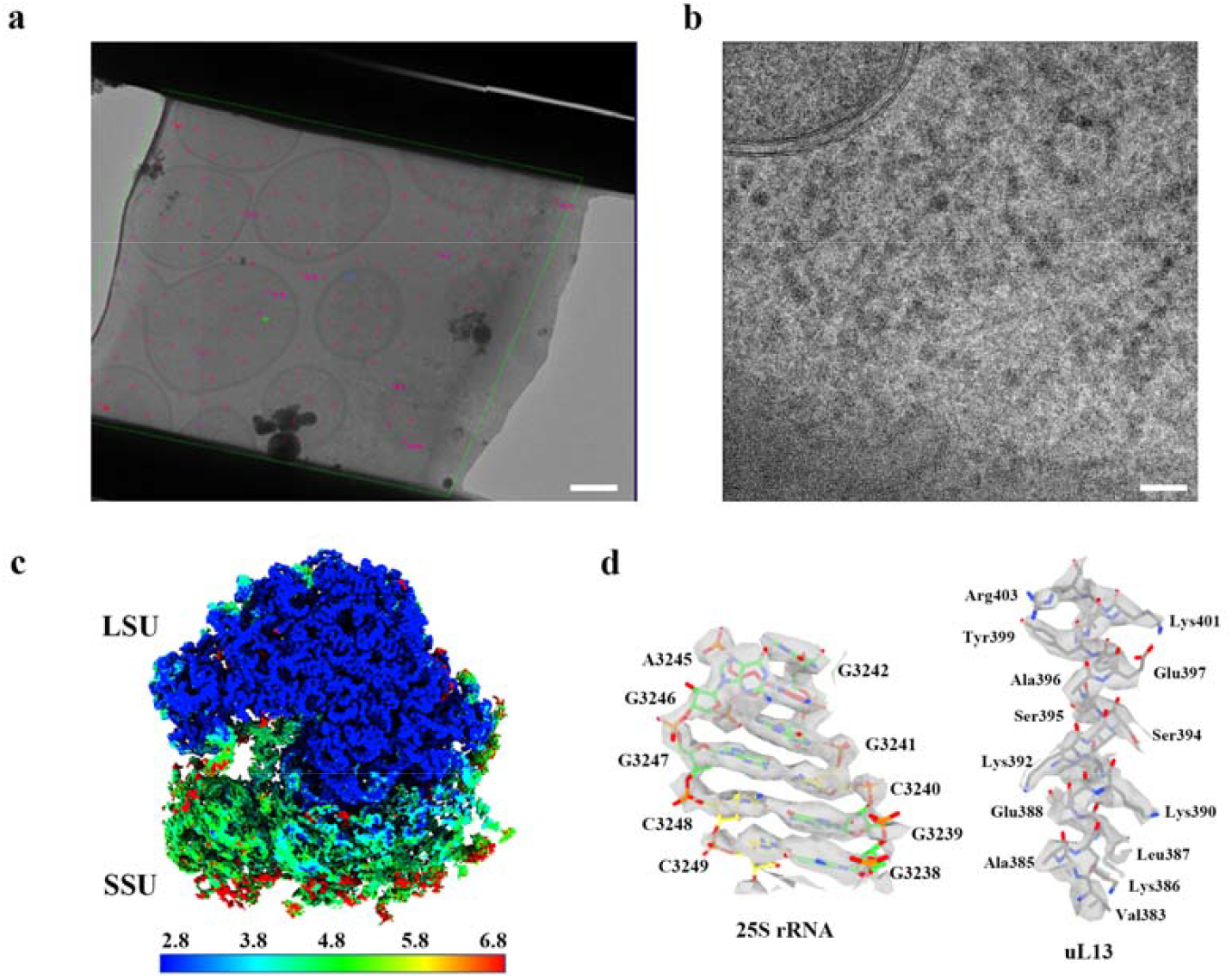
In situ *S. cerevisiae* ribosome structure. **a**, Overview of the cellular lamella. The magenta crosses indicate the data collection areas. The scale bar is 1 μm. **b**, A micrograph showing the cellular context of *S. cerevisiae*. The scale bar is 50 nm. **c**, Local resolution map of the averaged 80S ribosome. **d**, Well-resolved bases of 25S rRNA (left) and side chains of ribosomal protein uL13 (right).

**Fig 2.**
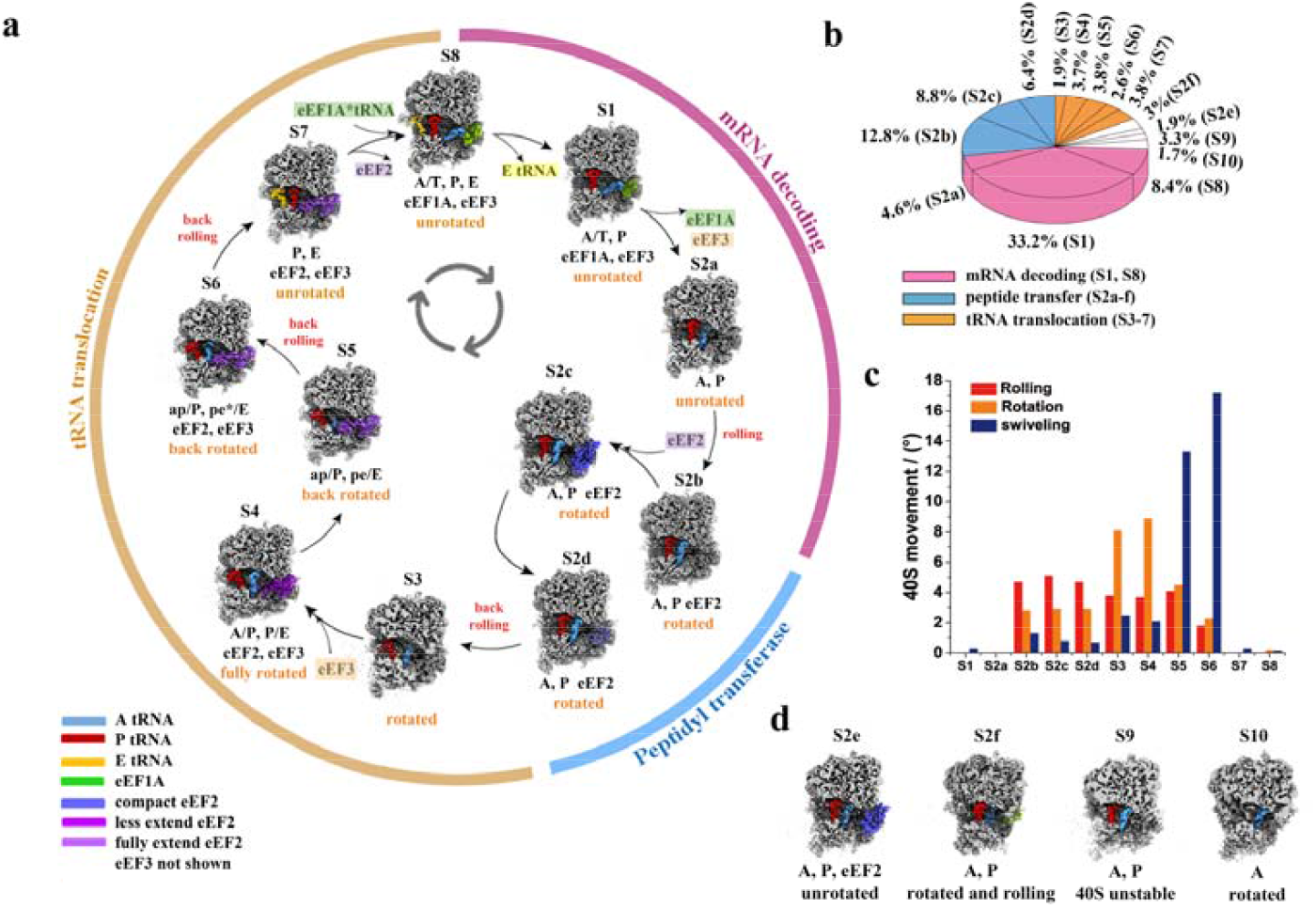
Ribosomal conformations and the translation elongation cycle. **a**, The elongation cycle is represented by eleven structures, distinguished by eEFs and tRNAs. These structures correspond to distinct phases: mRNA decoding (pink), peptidyl transferase (blue) and tRNA translocation (orange). The states of the 40S subunit rotation (orange) and rolling (red), along with the positions of tRNAs and elongation factors, are highlighted using different color shadings. **b**, The frequencies of occurrence for different states are calculated by the classified particle numbers. States corresponding to each process are colored according to (a). Four states unassigned in the elongation cycle colored in white. **c**, The angles of rotation (orange) and subunit rolling (red) for the 40S body and swiveling (blue) angles of the 40S head in eleven main conformational states, relative to S2a. **d**, Densities of four additional structures cannot be placed in the sequential cycle. Among these, state 10 contains only a tRNA in the A-site. The tRNAs and eEFs were colored the same as (a).

We reconstructed an intact translation elongation cycle of *S. cerevisiae* according to the assignment of the three eEFs (eEF1A, eEF2 and eEF3) and the locations of tRNAs binding to the ribosome. The elongation cycle consists of eleven main conformational states (Fig. 2a), and the corresponding percentage of particles accounted for by each state are listed in Fig. 2b and Supplementary Table 1, which is similar with previous studies in situ^6-8,12^. Of the eleven conformations (Fig. 2a), three states, state 8 (‘A/T, P, E’, S8), state 1 (‘A/T, P’, S1) and state 2a (‘A, P’, S2a), represent the ribosome undergoing the mRNA decoding process, collectively accounting for 46.2% of particles. Three ‘A, P’ states are newly resolved and present the stepwise native peptide transfer process (states 2b-2d, S2b-2d), comprising 28% of particles. Four classical states, state 4 (‘A/P, P/E’, S4), state 5 (‘ap/P, pe/E’, S5), state 6 (‘ap/P, pe*/E’, S6) and state 7 (‘P, E’, S7) and a new intermediate state of state 3 (S3) collectively accounting for 15.8% of particles, belong to tRNA translocation were obtained. The rotation of the 40S body and swiveling of the 40S head in each state were measured (see Methods, Fig. 2c and Supplementary Table 2). Three of these main conformations were further subdivided into nine sub-conformations according to the presence of eEF3 (Extended Data Fig. 2a, c, d), wherein a new state 4a (S4a), featuring the binding of eEF3 to 80S, was captured. Furthermore, four additional states may not be part of the classical cycle and are listed separately in Fig. 2d. The results were confirmed through two additional rounds of 3D classification (Supplementary Table 3).

### Compact eEF2 binds to the ribosome during peptidyl transfer

Classifications on particles in the “A, P” state by focusing on 40S subunit and eEF2 binding regions yielded seven different conformations. Among these, S2a belong to tRNA accommodation and three states (S2b-d) were associated with peptidyl transfer process. During peptide bond formation, the nascent peptide chain is transferred from the P-site tRNA to the newly incorporated A-site tRNA in the “A, P” state. Although 3D classification did not focus on the PTC, the resulting three states (S2b-d) represent the stepwise process of peptidyl transfer. States 2e (‘A, P’, S2e) and 2f (‘A, P’, S2f) were intentionally not included as main states of tRNA accommodation or peptidyl transfer process, considering the controversy surrounding the description of peptidyl chain transfer because of the relatively low resolution in PTC regions. The remaining state 9 (‘A, P’, S9), consisting of 3.3% of ribosome particles with heterogeneities in the 60S and 40S subunits (Fig. 2d and Supplementary Fig. 1), probably does not belong to the reconstructed elongation cycle.

In the unrotated S2a, tRNAs transition from “A/T, P” state to “A, P” state with the dissociation of eEF1A, associated with classical tRNA accommodation^5,13^. However, the density of the aminoacyl-tRNA elbow is poorer than that of the anticodon stem-loop and the CCA of the A-site tRNA is not observed in the PTC, indicating that the accommodation of A-tRNA is incomplete. The nascent peptide chain is attached to the P-site tRNA. The S2b exhibits a eukaryote-specific ‘subunit rolling’ of a ∼5° tilt relative to unrotated states, occurring towards the P stalk around the previously defined axis^14,15^ located on the upper part of h44 of 18S RNA and orthogonal to the well-known inter-subunit rotation axis. This rolling is earlier than the ‘subunit rolling’ observed in the PRE classical state, as reported previously^14-16^, and may facilitate the accommodation of aminoacyl-tRNA^5,14,15^. The CCA of A-site tRNA inserts into the PTC, and the nascent peptide chain is still connected to P-site tRNA in this state (Fig. 3a), indicating the pre-peptidyl transfer state upon accommodation of A-site tRNA. An intermediate state, S2c, during peptide bond formation was also captured (Fig. 3b, c). Peptide chain nucleophilic attack by the incoming free α-amino group onto the carbonyl carbon is known to break the acyl linkage between the carbonyl carbon and the 3’ hydroxyl of the P-site tRNA^17^ and form the peptide bond between the carbonyl carbon and the amino nitrogen. In S2c, the peptide chain is not connected to A-or P-site tRNA at a contour value of 2.6 RMSD (Fig. 3b). Decreasing the value to 1.8 RMSD revealed that the peptide chain connects to the two tRNAs simultaneously (Fig. 3c), suggesting that this intermediate represents the tetrahedral intermediate of the nucleophilic attack. The CCA of P-site tRNA is observed to interact in the PTC with G2620, G2621 and A2820 and that of A-site tRNA interacts with G2922, U2923, U2924 and U2953^17-19^ (Fig. 3d). Additionally, the density of the CCA of A-site tRNA is poorer than that in S2b, and the aminoacyl residues in S2c appears to shift slightly relative to S2b (Extended Data Fig. 3). In S2d, the peptide chain is well connected to A-site tRNA, representing the final peptidyl transfer state (Fig. 3e). The 40S subunit remains stationary and in the rolling state throughout peptide transfer process (S2b-d).

**Fig 3.**
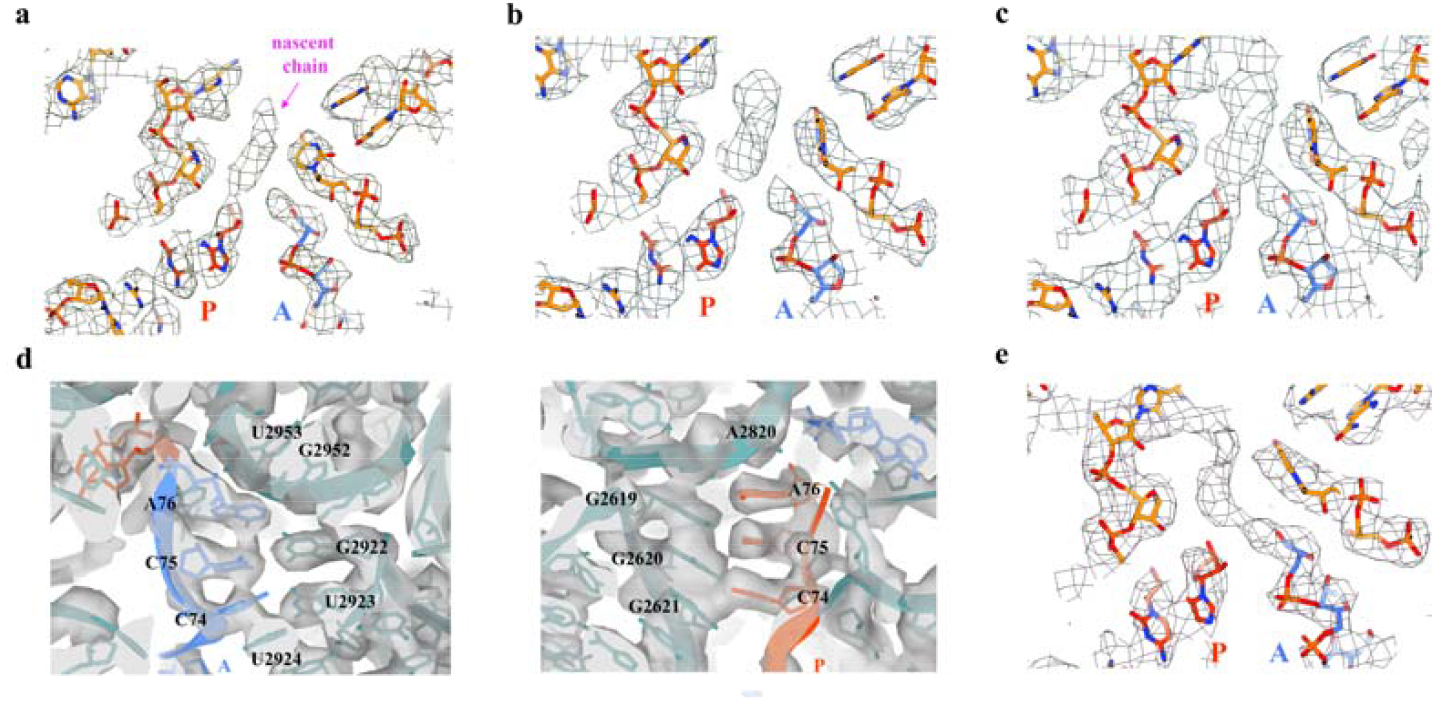
Presentations of cryo-EM densities in PTC during peptidyl transfer process. **a**, The positions of the nascent peptide chain, CCA of P-site tRNA (carbon atoms of model colored in red) and A-site tRNA (carbon atoms of model colored in blue) in PTC of S2b. **b-c**, View of the nucleophilic attack in S2c at a contour value of 2.6 RMSD (b) and 1.8 RMSD (c). **d**, Surroundings of CCA of A-site tRNA (left, model colored in blue) and P-site tRNA (right, model colored in orange) in S2c. Densities are colored in gray. **e**, The peptide chain disconnects with P-site tRNA (red) and is connected to A-site tRNA (blue) in S2d, indicating that the peptide transfer process is complete. The contour level is 2.6 RMSD in (a) and (e). Densities in (a), (b), (c) and (e) show in mesh.

Interestingly, besides transferring the nascent peptidyl chain in the PTC, we observed a clear occupation of eEF2 in S2c. When docking the five domains of eEF2 into the EM maps independently, as shown in Fig. 4a, b, we found that eEF2 exhibits a compact arrangement, which resembles the conformation of elongation factor G (EF-G) in prokaryotes^20^. Domains I and V connect with the 60S subunit, exhibiting a similarity to the typical extended conformation^21^. Domain II interacts with the 40S body through eS24, h15 and h17, whereas domain III contacts uS12 of the 40S body. Additionally, loop1 and loop3 of domain IV are proximal to the shoulder of SSU through h17 and h18, respectively. These observations indicate that compact eEF2 immobilizes the tilted SSU on LSU in S2c, where nucleophilic attack may occur, potentially ensuring a stable environment and optimal performance for peptide transfer. Compact eEF2 was also captured in unrotated S2e (Fig. 2d and Extended Data Fig. 4a, b), supporting observations from previous studies in prokaryotes^20^. Compared with eEF2 in S2c, we observed that in S2e, the interactions between domains II and IV of eEF2 and the 40S subunit are absent. eEF2 in S2b, S2d and S2f has a weak density, which may represent intermediate states of eEF2 undergoing conformation changes.

**Fig 4.**
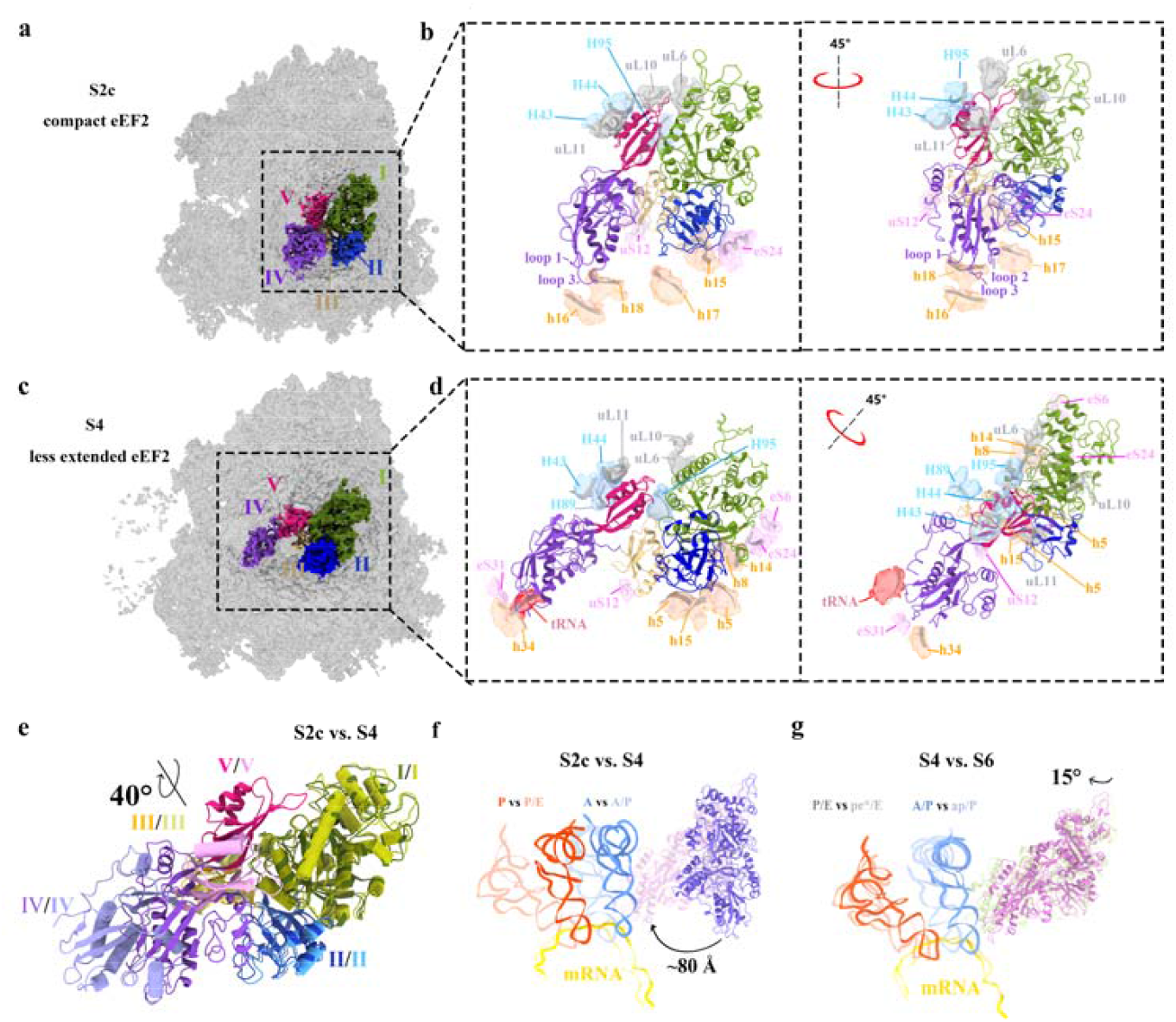
Compact and extended eEF2 during elongation cycle. **a**, The overview of the compact eEF2 bound to ribosomes in S2c. The cryo-EM densities of eEF2 are colored differently: domain I (green), domain II (blue), domain III (yellow), domain D (purple) and domain V (magenta). **b**, Two views of the environment surrounding of compact eEF2 (colored by domain) in the ribosome. Gray and blue densities are nucleic acids and proteins of LSU, respectively. Orange and pink densities are nucleic acids and proteins of SSU, respectively. The atomic models of the ribosome are shown in dark gray ribbon. Domain I of eEF2 interacts with uL6 and uL10 of the 60S subunit, and domain V interacts with H43, H44, H95 and uL11 of the 60S subunit. **c**, Less extended eEF2 in S4. **d**, Two views of the environment surrounding of less extended eEF2 in the ribosome. The arrangement and color are the same as (b), with the new densities of tRNA interacted with domain IV of eEF2 colored in red. **e**, Superimposing the models of compact and less extended eEF2. Helices in the less extended eEF2 are displayed by cartoon tubes. The domains colored same as (b) and (d), while less extended eEF2 are represented by a lighter color compared to compact eEF2. **f**, Overall movement of eEF2 and tRNAs on the ribosome from S2c to S4. The conformational change is shown from the compact (purple, in S2c) to the less extended conformation (pink, in S4) of eEF2. The A-, P-site tRNA and mRNA in S2c colored in blue, red and yellow, respectively. The translocated tRNAs and mRNA in S4 represented by light color. **g**, The conformational change from the less extended (pink, in S4) to fully extended conformation (light green, in S6) of eEF2. The tRNAs and mRNA in S6 represented by light colored compared to those in S4. The distances and angles presented reflect the movement of eEF2.

We also found clearly resolved densities belonging to eS25 (sequence 4-21) during the peptide transfer process in S2b-d (Extended Data Fig. 5c), which was not previously modeled. We used Alphafold2 to predict the eS25 structure, and sequence 4-21 was predicted to form an α-helix, aligning well with our cryo-EM densities. This helix interacts with A-site tRNA, extends over P-site tRNA (Extended Data Fig. 5c) and may stabilize aminoacyl-tRNA during peptidyl transfer. However, the helical densities in unrotated S2a and S2e exhibited poor continuity (Extended Data Fig. 5a, b) and were absent in other states beyond peptidyl transfer process.

### Conformational changes of eEF2 are coupled with translocation

After peptidyl transfer, peptidyl-tRNA begins translocation from the A-to P-site. S3-8, with the peptidyl chain connecting with A-site tRNA, represent intact eukaryotic ribosomal translocation and is well-sorted according to the positions of P- and E-site tRNA (Fig. 2a). eEF2 adopts two different conformations during translocation, the less extended and fully extended states. In S3, the SSU body undergoes a back rolling of ∼1° and a large relative rotation of ∼8° with respect to unrotated states. A weaker density of eEF2 compared with that in S2d and a flexible density of A-site tRNA (Supplementary Fig. 1) indicate that S3 represents an intermediate state at the beginning of translocation. Following another small SSU body rotation (Fig. 2c and Supplementary Table 2), a less extended conformation of eEF2 is observed in S4 (Fig. 4c, d), where domain IV does not overlap with the A-site on the 40S subunit, as previously resolved in situ^8,22^. The surroundings of less extended eEF2 are shown in Fig. 4d. Domain IV is primarily maintained by the presence of peptidyl-tRNA, with additional support from h34 and eS31. Domain III maintains its interactions with the SSU body through uS12, whereas domain I establishes interactions with 40S subunit through eS24, eS6, h8 and h14, facilitated by the rotation of the 40S subunit. Simultaneously, h5, near domain II, may stabilize eEF2 during its conformational change. As tRNAs move along the mRNA, the SSU head swivels ∼13.3°, and the SSU body undergoes a ∼3.6° back rotation in S5 (Fig. 2c and Supplementary Table 2). eEF2 is found to extend towards the A-site and reaches the fully extended state^21^. The SSU body continues with another small back rotation of ∼2° while the head swivels to ∼17.2°, leading to tRNAs reaching S6 with a fully extended eEF2^16^. The interactions between domains I and II of eEF2 with SSU are mostly lost with the back rotation of the SSU body (Extended Data Fig. 4d). A back rolling of ∼2.1° was observed between S5 and S6, which may, together with the back rotation, facilitate tRNA-mRNA decoupling between the SSU body and head^16,21^. In S7^7^, the SSU body returns to the unrotated and unrolling state while the head swivels to 0°. Simultaneously, peptidyl-tRNA reaches the P-site, and deacyl-tRNA occupies the E-site, indicating that the translocation process is nearing completion. eEF2 maintains a fully extended conformation until dissociation in S8. Another elongation cycle begins when eEF2 separates, and eEF1A·tRNA combines with the next mRNA codon.

We compared the domain arrangement of compact eEF2 in S2c and less extended eEF2 in S4 by superimposing their models (Fig. 4e), revealing that domain I-III maintain a relatively fixed arrangement among each other, and domains IV-V form another rigid body, undergoing a rotation of ∼40° around an axis at their junction. This rotation causes domain IV of compact eEF2 to extend about 48 Å, mostly towards the A site of the 40S subunit, whereas domain V shifts ∼23 Å to be primarily closer to 40S subunit (Fig. 4e). The spatial changes of eEF2 resulting from conformational changes are illustrated using the 60S subunit as a reference to analyze the shifts and rotations of eEF2 domains relative to the ribosome. From compact to less extended, eEF2 domain I-III rotated ∼30° and domain IV-V rotated ∼60°, yielding a movement of ∼80 Å for domain IV (Fig. 4f). From less extended to fully extended, eEF2 domains have a monolithic rotation of ∼15°, leading to domain IV of S6 being extended primarily by ∼20 Å towards the A-site (Fig. 4g).

Conformational changes of eEF2 from the compact state in S2c to the fully extended state in S6 do not affect the interactions of domain V with H95 of 25S rRNA, H43 and H44 of the P stalk (Extended Data Fig. 6). By aligning domain V of fully extended eEF2 with compact eEF2 on the ribosome in S2c, we observed steric hindrance caused by the P stalk. This results in a clash of domain I with the 60S subunit (Extended Data Fig. 7a), hindering the formation of extended eEF2. When the ribosome transitions to S4, the P stalk shifts ∼7 Å towards tRNA (Extended Data Fig. 7b), and the 40S body rotates by ∼6° to allow eEF2 to adopt the extended conformation. We found a stronger density of uL10 of the P stalk near domain I when focusing on the density of eEF2 in S3 (Supplementary Fig. 2).

### Conformational rearrangement of eEF3 throughout elongation

In addition to the two conserved and shared elongation factors, eEF1A and eEF2, found in eukaryotes, there is a requirement for another ATPase known as eEF3 in the fungal translation system^23-25^. This ATPase contains two ATP-binding motifs and spans the head of the 40S to 60S subunits. Focus on the overall cryo-EM maps of S4, S8 and S1, the eEF3 essentially exhibit weak density, which may suggest a relatively low occupancy of eEF3. Therefore, a dedicated round of 3D classification focusing on the eEF3 region illustrated the 3D variability of eEF3 in these three states. S4 is divided into two sub-states, S4a (with stable eEF3 binding) and state 4b (S4b, without eEF3 binding), as illustrated in Extended Data Fig. 2a. Three sub-states are identified in S8 (Extended Data Fig. 2c): stably bound eEF3 (state 8a, S8a), flexibly bound eEF3 (state 8b, S8b) and unbound eEF3 (state 8c, S8c). Four sub-states are identified in S1 (Extended Data Fig. 2d): the stable binding to 80S state (state 1a, S1a), the less-flexible binding state (state 1b, S1b), the more-flexible binding state (state 1c, S1c) and the dissociated state (state 1d, S1d), where eEF3 may dissociate from the 80S.

Two conformations of eEF3 were identified from eEF3-binding states. A stable closed-eEF3 conformation (Fig. 5a) when bound to the unrotated ribosome in S1a and S8a (Extended Data Fig. 2c,d), which closely resembles the state obtained from single-particle analysis^24^, and a stable open-eEF3 conformation (Fig. 5b and Extended Data Fig. 2a) when bound to the fully rotated ribosome of S4a, which has not been observed previously. The local resolution of open-eEF3 is lower than closed-eEF3 (Supplementary Fig. 3) because of inherent flexibility. When docking the eEF3 models into the EM density, two ATP-binding domains (ABC1 and ABC2) of open-eEF3 (Fig. 5b) exhibit significant gaps between the HEAT domain and the ribosome compared with those of closed-eEF3 (Fig. 5a). The analysis of chromodomain (CD) is absent in our structures given that the density of CD is insufficient for model fitting. An overarching perspective shows that the closed-eEF3 exhibited a pronounced rotation relative to open-eEF3 around the junction of the ABC2 domain and the ribosome, primarily attributed to the rotation of the 40S subunit (Fig. 5c). When the HEAT domain and four-helix bundle domain (4HB domains) are superimposed, ABC1 and ABC2 domains, which are in complex with ATP or ADP, have a rotation of ∼90° on the axis of the HEAT domain (Fig. 5d).

**Fig 5.**
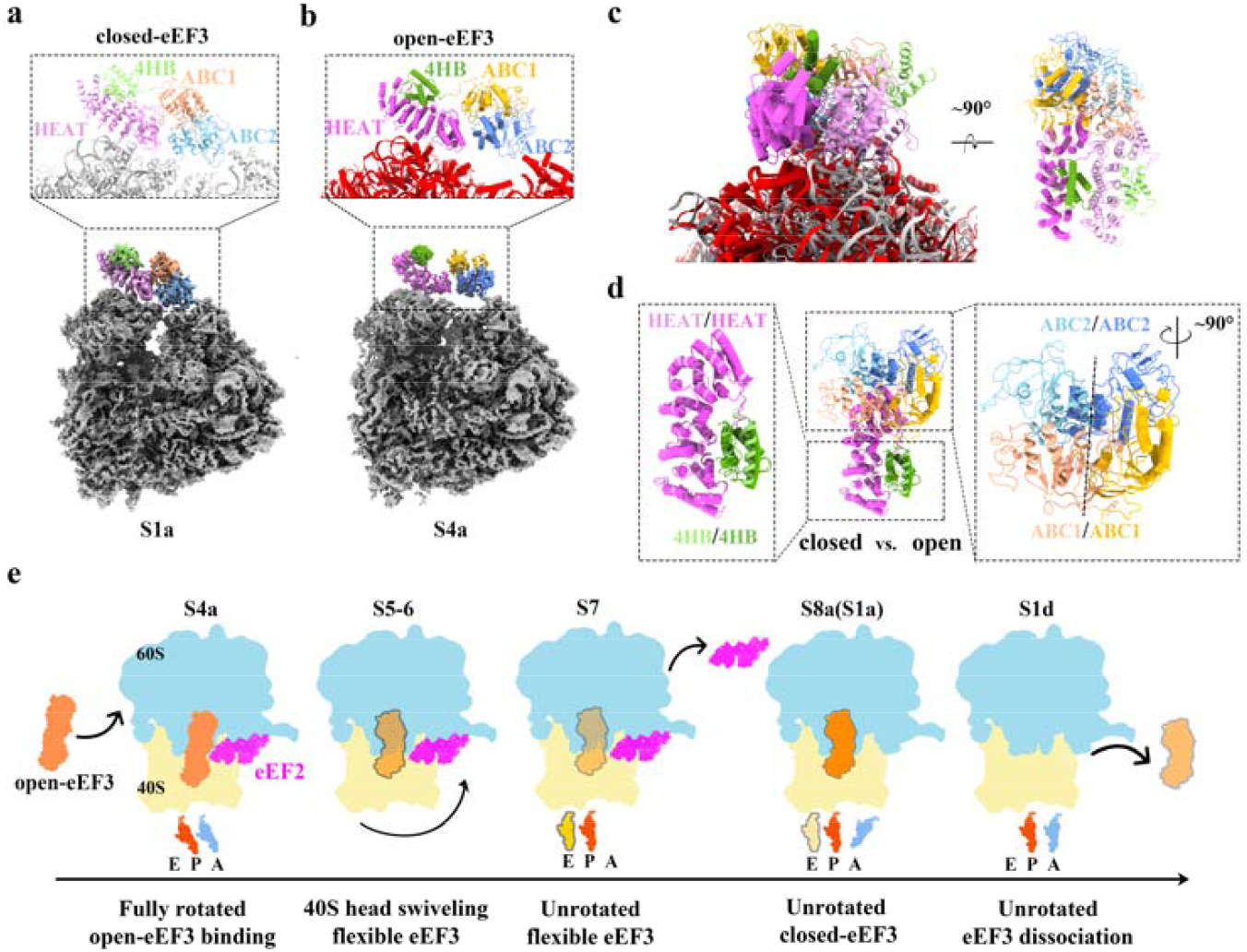
eEF3 binds to and dissociates from the ribosome in different conformations. **a**, Bottom1 panel is the overall densities of closed-eEF3 in complex with the unrotated ribosome (S1a). Detailed zoom highlighting eEF3 domains: HEAT (pink), 4HB (green), ABC1 (orange) and ABC2 (blue). Top panel show close-up of model located in closed-eEF3 region. The eEF3 domains colored consistently with their densities, while model of ribosome colored in gray. **b**, The presentation of open-eEF3 in complex with the fully-rotated ribosome (S4a). The domains of eEF3 colored by a darker color compared to that of the closed-eEF3 in (a), with helical models are displayed by a cartoon tube. The model of rRNAs colored in red. **c**, Comparison of closed-eEF3 and open-eEF3 under superimposition of LSU models of S1a and S4a. **d**, Swinging between the ABC1 and ABC2 domains of the two eEF3 conformations under superimposition of HEAT and 4HB domains. The colors are same as (a). **e**, Binding states of eEF3 vary with the ribosome states. Stable and flexible binding eEF3 are illustrated in orange and light orange, respectively. The intermediate color between orange and light orange in the second pattern represents the occurrence of closed eEF3 in S6. The states of tRNAs and SSU were indicated below the corresponding ribosomes. The tRNA colored by light yellow in the fourth pattern represents the release of E-site tRNA in S1a. The purple outline represents eEF2 in the elongation cycle.

Throughout the elongation cycle, the open-eEF3 initially binds to the ribosome when the 40S subunit reaches the fully rotated state of S4a (Fig. 5e and Extended Data Fig. 2a). As the 40S subunit undergoes body back-rotation and large head swiveling, the stability of eEF3 binding to the ribosome weakens, leading to flexible eEF3 density in S5 and subsequently transitioning to a closed conformation in S6 (Extended Data Figs. 2b and 8). However, this quality of eEF3 density in S6 is lower than that of the closed-eEF3 observed in S1a. The flexible density of eEF3 in S5 indicates a conformational change of eEF3 coupled with head swiveling and back rotation of ribosomal SSU (Fig. 5e). Accompanying the head swiveling of the 40S subunit from S4 to S6, the E-site tRNA is transferred ∼14 Å and the L1 stalk opens by ∼20 Å in the presence of the 40S body alignment (Extended Data Fig. 9a).

Following the dissociation of extended eEF2 and the binding of the eEF1A·tRNA complex in S7, strong densities are observed for eEF3 in the closed conformation in S8a and S1a. In S8a, the L1 stalk continues to open ∼19 Å relative to S6, facilitating the release of the E-site tRNA from the ribosome. The L1 stalk adopts open conformation in three sub-states identified from S8. After releasing the E-site tRNA in S1 (Fig. 2a), the L1 stalk of the 60S subunit switches to the closed state. In the stable eEF3-binding state of S1a, the L1 stalk of 80S adopted the open state, whereas, in the dissociated state of S1d, L1 switched to the closed state. To verify whether the transition of L1 stalk from open to closed is sequential, we classified particles belonging to ‘A/T, P’ state (S1) into three classes according to the state of the L1 stalk: open state, intermediate state and closed state (Extended Data Fig. 9b) with the percentage of the stable eEF3 binding state calculated as 73%, 26% and 0%, respectively. From the results, we infer that the presence of E-site tRNA does not affect the binding state of eEF3, and the closure of the L1 stalk must occur after eEF3 dissociation.

## Discussion

In this study, we resolved the dynamic structures of the S. cerevisiae ribosome in a native translation environment at near-atomic resolutions. By following tRNAs and eEFs, we gained insight into the life cycle of ribosomal elongation. The compact conformation of eEF2 during peptidyl transfer process was captured, which occurs much earlier than previous observed extended eEF2^20,26^. In addition, we found an open-eEF3 conformation binding to fully rotated ribosome, whose conformational change may facilitate the head swiveling of 40S subunit.

Subunit rolling is a common phenomenon of SSU in eukaryotic peptidyl transfer, which facilitates the accommodation of aminoacyl-tRNA^14,15^. In our findings, rolling SSU was fixed by compact eEF2 to the LSU in S2c, where the peptide chain undergoes nucleophilic attack by the free α-amino. It is obvious that compact eEF2 binds to the ribosome at or before S2c. The absence of CCA of A-site tRNA in PTC of unrotated ribosome indicates uncompleted accommodation. Therefore, we hypothesize that the unrotated S2e, characterized by compact eEF2 binding and previously unassigned in the elongation cycle, may belong to A-site tRNA accommodation given its unrotated SSU and absence of CCA of A-site tRNA in the PTC. The helix of eS25 exhibits lower densities in S2a and S2e compared to that in S2b-d (Extended Data Fig. 5), suggesting that S2e is a state close to S2a. In addition, subunit rolling reduces the openings to the inter-subunit space on the A-site side, which does not facilitate eEF2 binding in S2c.

Here, we present a model (Extended Data Fig. 10) to provide insight into the life cycle of compact eEF2 during elongation cycle. Following the dissociation of eEF1A and eEF3, the aminoacyl-tRNA approaches to the A-site, but the CCA does not insert into the PTC. In S2a, unrotated ribosomes serve as substrates for compact eEF2 in the cytoplasm and facilitate ribosome recruitment. The compact eEF2 binds only to 60S subunit in S2e because the eEF2 can’t reach SSU body in unrotated ribosome. Upon eEF2 binding, the ribosome undergoes a conformational change, tilting the 40S subunit by ∼5° towards the P stalk, and attempts to establish an interaction with eEF2 that gives rise to a flexible density of eEF2 in S2b. When eEF2 establishes a stable interaction with both 60S and 40S subunits simultaneously, the CCA is inserted into the PTC, and transpeptidation proceeds stably with the assistance of eEF2 in S2c. As the peptide transfer is completed, the SSU prepares for back-rolling and rotation to enable the conformational change of eEF2 and eEF2 exhibits weak density again in S2d. A compact EF-G binding to the unrotated 70S ribosome before peptidyl transfer in *E. coli* was captured previously by taking advantage of nonhydrolyzable aminoacyl-tRNA analogs in P-site tRNA^20^, which supports our hypothesis. Recently, Hassan et al. showed that subunit rolling is common in prokaryotes and eukaryotes^27^, suggesting that compact eEF2 or EF-G in the ribosomal elongation cycle is ubiquitous.

After analyzing different structures of eEF2 at different elongation states, our results indicated that eEF2 play a pivotal role as a ‘molecular arm’ in the eukaryotic elongation cycle. Specifically, during peptidyl transfer process, the loops of domain IV of compact eEF2 function as ‘fingers’ within the contracted arm, skillfully captures the tilted SSU body and facilitates the seamless transfer of the nascent peptide. As the cycle progresses to translocation, the ‘arm’ gradually extends to reach the A-site. Simultaneously, the ‘fingers’ play a crucial role in stabilizing the mRNA reading frame, effectively ensuring the directionality of the translocation process^21,28^, which corresponds to states S5-7 in our results.eEF2 is a GTP-binding protein^29-31^ and plays crucial roles during the ribosomal elongation cycle. Majumdar et al.^16^ resolved an extended eEF2·GDP binding ribosome in *Giardia intestinalis*, featuring a uniquely positioned ‘leaving phosphate’. This structure aligns with our S6 state, as confirmed by comparing tRNA positions with SSU body alignment, thus revealing that GTP hydrolysis occurs during the final steps of tRNA translocation^16,28^. In addition, we hypothesize that the large SSU rotation at the beginning of translocation (S3 and S4) induces the conformational change of eEF2, and this change does not require GTP hydrolysis by eEF2. The small back rolling of ∼1° from S2d to S3 weakens the interaction between domains II and IV of eEF2 and the SSU body, facilitating the subsequent extensive body rotation in S3. However, the interaction between domain III and uS12 of 18S rRNA is maintained, accompanying the SSU rotation and dragging the compact eEF2 in rotation around the junction of domain I and 60S subunit. Simultaneously, the closure of H95 in 25S rRNA removes steric barriers for extending eEF2. The uL10 of the P stalk likely plays a key role in facilitating the movement of domain V (Supplementary Fig. 2).

Hydrolysis of GTP (ATP for eEF3) occurs when the elongation factors maintain ribosome translation, along with conformational changes depending on whether they are bound to GTP (ATP for eEF3) or GDP (ADP for eEF3). eEF3 contains two ATP-binding domains, thus carrying two ATP molecules^24,25^. We hypothesize that the open-eEF3 with two ATPs binds to the ribosome in the fully rotated ribosome of S4a. One ATP hydrolysis occurs during the conformational change of eEF3 from open (S4a) to closed (S6) states and may facilitate head swiveling and E-site tRNA translocation to exiting. In the S4, the E-site tRNA binds to the L1 stalk, and the translocation of tRNA towards the L1 stalk potentially drives the opening of the L1 stalk, facilitating the release of the E-site tRNA. We hypothesize that the opening of the L1 stalk and release of E-site tRNA occur without additional ATP hydrolysis by eEF3. The second ATP hydrolysis may contribute to dissociating eEF3 from the ribosome, where eEF3 binding is destabilized before dissociating in S8. If this occurs, the second ATP hydrolysis causes a conformational change of eEF3 that disrupts the interaction interface between eEF3 and the ribosome to cause eEF3 dissociation from the ribosome.

We identified 451,700 true positive particles based on the presence of the SSU, which was not included in our template. Notably, the tRNAs and eEFs analyzed in the elongation cycle were also not included in the template. Additionally, the frequencies utilized for detection by GisSPA ranged from 1/400 Å^−1^ to 1/8 Å^−1^. Consequently, the high-resolution signals above 1/8 Å^−1^ in the ribosome remain unaffected by template bias. Furthermore, following the final 3D classification, the densities of the LSU included in the template and the SSU beyond the template exhibited equivalent quality in each stable state. These comprehensive analyses collectively indicate that our results are hardly affected by model bias. In addition, we did not discuss the distribution of each state within the cell. Although the Z coordinates of our ribosomal particles can obtained from defocus searching, similar to the 2DTM of Lucas et al.^32^, the precision is significantly lower than that of cryo-ET and is inadequate for accurately repositioning 20-nm sized ribosomes in a cellular context, as discussed in our previous studies^9^.

The single LSU and ribosomal conformations in the initiation and termination states were not observed even though we have successfully identified conformations at a percentage of 1.5%. Thus, the absent states in our study present lower percentages than 1.5% in *S. cerevisiae* cells. We obtained 451,700 ribosomes after collecting data for 2 days. The large dataset increased the resolution of the ribosome at each state and facilitated the capture of an intermediate state with an estimated occupancy as low as ∼1%. The use of GisSPA provide significant advantages in terms of high data acquisition throughput.

## Methods

### Cell culture

Saccharomyces cerevisiae strains BY4741 were grown in YPD medium at 30 °C to an OD600 of 0.6 ∼ 0.8. The cells were collected and resuspended in the same medium, and the concentration of the suspension was adjusted to an OD600 of 2.

### Cryo-EM sample preparation and FIB milling

3 µL *S. cerevisiae* cells were applied to a glow-discharged copper Quantifoil 200 mesh R1.2/1.3 grid. After 30s pre-blotting the grid was plunge-frozen into liquid ethane using a Leica EM GP (Leica Microsystems) with 6 ∼ 8 s back-side blotting at 16 □ with the chamber at 70% humidity.

Cryo-lamellae were prepared by cryo-FIB milling using the Aquilos-2 FIB-SEM microscope (Thermo Fisher Scientific). Before milling, samples were sputter-coated with metallic platinum for 10s with 15 mA beam current then coated with organo-platinum for 6s through the gas-injection-system (GIS) and coated metallic platinum for 10s again with the same settings. Lamellae were rough milled to ∼300 nm using a decreasing Ga+ ion beam from 0.3 nA to 50 pA with a milling angle of 8°. 10 pA ion beam was used for the final polish to obtain a final ∼ 150 nm thickness.

### Cryo-EM data acquisition and image processing

Approximately 40 cryo-lamellae were collected by applying a single particles data collection method^33^ using FEI Titan Krios G3 (Thermo Fisher Scientific) equipped with a Gatan GIF K2 camera (Gatan, Icn.). Before collection, an offset start angle of 8° was set to compensate for the milling angle, ensuring that the lamellae perpendicular to the electron beam details refer to previous description^11,34^. Data collection was performed with SerialEM^16^ and the beam-image shift data collection method^17^ using a nominal magnification of 105,000 with a calibrated pixel size of 0.66 Å/pixel in super-resolution mode.

Two sets of dose rate were used during data collection, 219 micrographs were irradiated by 35 e^-^ and dose-fractionated to 40 frames, and the rest were collected with a total dose of 50 e^-^ and dose-fractionated to 50 frames. The energy slit width was set to 20 eV and the nominal defocus was set between -0.8 to -1.2 μm.

### Structure determination

Motion correction was done by MotionCor2^35^ and micrographs were dose-weighted and binned with a factor of 2. The CTF parameters of micrographs were estimated by CTFfind4^36^ in RELION, then the micrograph file in “star” format was converted to “lst” format for the detection task. The 60S subunit structure of yeast (EMD-6615)^37^ was used as the initial 3D model in GisSPA^11^, which is relatively more homogeneous than the 40S subunit, as the initial 3D template. The localization task was performed by generating 1,649 2D projections (ignoring the in-plane rotation) that uniformly covered the sphere model with a 5° angular step. We performed the initial detection task on the first 500 micrographs with overlapping parameter *k* as 3 and output score threshold as 6.8, resulting in 7043 particles in total. A round of focused refinement by RELION^38^ on these particles yielded a structure of 60S at a resolution of 5.9 Å.

Considering the scale difference between the original template (EMD-6615) and the particles in our micrographs, which influenced the detection efficiency of GisSPA, we segmented the 60S subunit from the newly reconstructed map at 5.9 Å resolution and used it as the template for a new round of searching by GisSPA. To remove duplicated detections, potential particles within translational distance of ∼10.6 angstroms and Euler distance of 8° were merged as one particle. Only frequencies from 1/400 Å^−1^ to 1/8 Å^−1^ were used in the calculations for both localization tasks. 808,736 potential particles from the 3,702 micrographs were kept after removing repeats that met the new cutting threshold of 6.2. The particle file in “lst” format was converted back to “star” format for particle extraction in RELION. Particles were 2-binned extracted with a boxsize of 200, after implementing a round of 3D classification using the 3D model in detection as the initial model and the lowpass filter was set to 20 Å, ∼56% of the particles including the 40S subunit density were selected and the rest showing bias features were thrown out (Extended Data Fig.1). Since the cryo-EM data on each grid was collected individually, we divided them in to 9 optical groups for the following refinements. 2 rounds of CTF parameters refinements, 1 round of beam tilt estimation and 2 rounds of orientational and translational refinements were performed, the resolution was estimated as 3.0 Å in 60S region. To further improve the resolution, a round of Bayesian polishing was carried out, and another round of focused refinement was applied to the polished particles, and the resolution in 60S subunit was improved to 2.9 Å. The distributions of the final selected particles in several representative micrographs are depicted in Supplementary Fig. 4.

### 3D classification and refinement

Ribosome 3D classifications were accomplished by CryoSPARC^39^ with the imported refined particles from RELION3.2. First, focused classification with a smooth shape mask covering the A-, P-, E-site tRNA positions was performed. This resulted in four major classes with different tRNA occupancies: “A/T, P” (150,024 particles), “A, P” (201280 particles), “A/T, P, E” (37,868 particles) and “P, E” (62,528 particles). In the second round, we continued to classify particles belonging to “P, E’ class focused on 40S region, and this resulted in five tRNA translocation classes divided from “‘P’, ‘E’” class: “A/P, P/E” (S4), “ap/P, pe/E” (S5), “ap/P, pe^*^/E” (S6) and “P, E” (S7) (Extended Data Fig.1).

We found densities with lower occupancy presented in the P stalk binding site in the “A, P” state, so a third round of classification focused on the P stalk binding site was carried out on particles in “A, P” state, resulting in empty binding site class, compact eEF2 stable binding class and compact eEF2 density scattered class. Each class showed flexibilities in 40S subunit, therefore, a classification focused on 40S region was performed on each class. Particles were further classified into the following states: unrotated 40S without compact eEF2 binding (S2a), rotated 40S without compact eEF2 binding (S2b), rotated 40S with compact eEF2 stable binding (S2c), rotated 40S with flexible compact eEF2 density (S2d), unrotated 40S with stable compact eEF2 binding (S2e), an intermediate state (state 2f) located between S2a and S2b and an intermediate class (S3) positioned between “A, P” and “A/P, P/E” states. In addition, 14,925 particles (S9) belonging to “A, P” state had a lower resolution of 4.8 Å than other classes within similar particles, showing severe flexibilities in 40S subunit, and 7,588 particles only contain a tRNA in A-site (S10) with 40S subunit partially rotated similar to S2b and S2c.

To investigate the occupancies of eEF3 in S1, S4 and S8, three rounds classifications with a soft mask focus on eEF3 region were individually performed on the particles in three classes, resulting in stable eEF3 (S1a, S4a and S8a), flexible eEF3 (S1b, S1c and S8b) and smeared eEF3 (S1d, S4b and S8c). Classifications of particles in “A/T, P” state focused on L1 stalk were performed with a soft mask including regions of L1 stalk.

The masks used above were generated by *relion_mask_creation* command implemented in RELION^38^, and the input maps were segmented from the overall reconstruction in Chimera^40^ and low-passed to 20 Å. The resolutions of all states were obtained through the final refinement focused on the 60S subunit. The local resolution calculated from CryoSPARC^39^.

### Determination of SSU removements

The movement analyses of the 40S subunit were conducted using Chimera^40^. The rotational angles, axes, and matrices of the 40S body between any state and S2a were obtained by aligning the 60S subunit of all maps. A custom script was employed to decompose the corresponding rotational angles along the axes of 40S body rotation and rolling. The rotational axis between S4 and S5 was chosen as the axis of 40S rotation, while the axis of subunit rolling was aligned with the upper part of h44 of 18S rRNA, identified in Chimera. Additionally, the angles of 40S head swiveling were calculated by Chimera through the alignment of the 40S body between each state and S2a.

### Model building

The *S. cerevisiae* 60S subunit model was built based on the crystal structure (PDB:4V88)^41^. It is split into separate rRNA and protein parts and fit into the cryo-EM density. The non-rotated 40S subunit part was built based on the PDB 7OSM^21^. For rotated 40S, PDB 7B7D^42^ was used. In addition, the refinement of PTC in 25S rRNA model refers to the 2.4 Å crystal structure (PDB :8CVJ)^4^. Alphafold2^43^ was used to complete the eS25 structure.

The eEF1A model was derived from the cryo-EM structure (PDB: 5LZS)^44^. Using PDB 2P8W^30^ as a template to build the elongated conformation of eEF2, the compact conformation of eEF2 needs to rigid-body-fit the five domains divided as previously described^45^ (domain I \ domain II \ domain III \ domain IV \ domain V) into the cryo-EM map by UCSF Chimera^40^. Another model (PDB:1N0V)^45^ is used for the rearrangement of the eEF2 compact conformation. The previous eEF3 model (PDB: 7B7D) was separated into five domains (HEAT \ 4HB \ ABC1 \ ABC2 \ CD)^2^ that were fitted to cryo-EM map separately as rigid body in Chimera. The tRNA comes from (7B7D), and the aminoacyl-tRNA model was derived from PDB ID 8CVJ. The mRNA was built using PDB 6XHW^46^ as template. The nascent peptide was built de novo by constructing a poly-alanine sequence.

These domains fitted into each electron density map as rigid bodies in Chimera^40^ via ‘fit in map’ function. Then manual adjust separate domains by real space refine and merge into a whole model in Coot^47^. The ribosome model is further refined by iterating in Coot and Phenix^48^. Validation was obtained using MolProbity7^49^.

## Data and code availability

The electron density maps have been deposited to the Electron Microscopy Data Bank (EMDB): EMD-38654(S8a), EMD-38655(S8), EMD-38656(S1), EMD-38657(S1a), EMD-38658(S2a), EMD-38659(S2b), EMD-38660(S2c), EMD-38661(S2d), EMD-38662(S2e), EMD-38663(S2f), EMD-38664(S3), EMD-38665(S4), EMD-38666(S4a), EMD-38667(S5), EMD-38668(S6), EMD-38669(S7), EMD-38670(S9), EMD-38671(S10), which are publicly available as of the date of publication. The atomic models have been deposited to the Protein Data Bank (PDB): 8Z71 (S1a), 8XU8 (S2c), 8YLD (S4a), 8YLR (S6), which are publicly available as of the date of publication.

The custom script of determination of SSU removements has been uploaded to GitHub: https://github.com/zhao-xigua/Rotation.

## Acknowledgments

We thank L. Kong for cryo-EM data storage and backup. The project was funded by the National Natural Science Foundation of China (32150010, 32325027, 31930069 and 32241027), the Strategic Priority Research Program of the Chinese Academy of Sciences (XDB37040101), the National Key R&D Program of China (2021YFA1301501), the Key Research Program of Frontier Sciences at the Chinese Academy of Sciences (ZDBS-LY-SM003) and Basic Research Program Based on Major Scientific Infrastructures, CAS (JZHKYPT-2021-05). Key Laboratory of Biomacromolecules, Chinese Academy of Sciences (ZGD-2023-05).

## Contributions

X.-Z.Z. supervised the project. C.-L.W. and J.-X.L. prepared the *Saccharomyces cerevisiae* cells, milled the cells to cellular lamellae by cryo-FIB and collected the cryo-EM Data. J.C. performed the data processing and conducted the analysis. J.-X.L. conducted the model buildings. X.-Z.Z., J.C., C.-L.W. and J.-X.L. wrote the initial draft, and all authors contributed to the discussion of the results and revision of the manuscript.

## Ethics declarations

### Competing interests

The authors declare no competing interests.

## Notes

### Competing Interest Statement

The authors have declared no competing interest.

### Summary of Updates

New manuscript has been revised to address the reviewers' concerns and some improper descriptions were also corrected.

## References

1. Dunstone, M.A. & de Marco, A. Cryo-electron tomography: an ideal method to study membrane-associated proteins. Philosophical Transactions of the Royal Society B: Biological Sciences 372, 20160210 (2017).

2. Andersen, C.B. et al. Structure of eEF3 and the mechanism of transfer RNA release from the E-site. Nature 443, 663–668 (2006).

3. Korostelev, A.A. The structural dynamics of translation. Annual Review of Biochemistry 91, 245–267 (2022).

4. Syroegin, E.A., Aleksandrova, E.V. & Polikanov, Y.S. Insights into the ribosome function from the structures of non-arrested ribosome–nascent chain complexes. Nature Chemistry 15, 143–153 (2023).

5. Loveland, A.B., Demo, G. & Korostelev, A.A. Cryo-EM of elongating ribosome with EF-Tu• GTP elucidates tRNA proofreading. Nature 584, 640–645 (2020).

6. Gemmer, M. et al. Visualization of translation and protein biogenesis at the ER membrane. Nature 614, 160–167 (2023).

7. Hoffmann, P.C. et al. Structures of the eukaryotic ribosome and its translational states in situ. Nature Communications 13, 7435 (2022).

8. Xue, L. et al. Visualizing translation dynamics at atomic detail inside a bacterial cell. Nature 610, 205–211 (2022).

9. Cheng, J., Li, B., Si, L. & Zhang, X. Determining structures in a native environment using single-particle cryoelectron microscopy images. The Innovation 2 (2021).

10. Cheng, J. & Zhang, X. Optimizing weighting functions for cryo-electron microscopy. Biophysics Reports 7, 152 (2021).

11. Cheng, J. et al. Determining protein structures in cellular lamella at pseudo-atomic resolution by GisSPA. Nature Communications 14, 1282 (2023).

12. Xing, H. et al. Translation dynamics in human cells visualized at high resolution reveal cancer drug action. Science 381, 70–75 (2023).

13. Rodnina, M.V. Decoding and recoding of mRNA sequences by the ribosome. Annual Review of Biophysics 52, 161–182 (2023).

14. Budkevich, T.V. et al. Regulation of the mammalian elongation cycle by subunit rolling: a eukaryotic-specific ribosome rearrangement. Cell 158, 121–131 (2014).

15. Behrmann, E. et al. Structural snapshots of actively translating human ribosomes. Cell 161, 845–857 (2015).

16. Majumdar, S. et al. Insights into translocation mechanism and ribosome evolution from cryo-EM structures of translocation intermediates of Giardia intestinalis. Nucleic Acids Research 51, 3436–3451 (2023).

17. Tirumalai, M.R., Rivas, M., Tran, Q. & Fox, G.E. The peptidyl transferase center: a window to the past. Microbiology and Molecular Biology Reviews 85, e00104–21 (2021).

18. Martin Schmeing, T., Huang, K.S., Strobel, S.A. & Steitz, T.A. An induced-fit mechanism to promote peptide bond formation and exclude hydrolysis of peptidyl-tRNA. Nature 438, 520–524 (2005).

19. Lapeyre, B. & Purushothaman, S.K. Spb1p-directed formation of Gm2922 in the ribosome catalytic center occurs at a late processing stage. Molecular cell 16, 663–669 (2004).

20. Lin, J., Gagnon, M.G., Bulkley, D. & Steitz, T.A. Conformational changes of elongation factor G on the ribosome during tRNA translocation. Cell 160, 219–227 (2015).

21. Djumagulov, M. et al. Accuracy mechanism of eukaryotic ribosome translocation. Nature 600, 543–546 (2021).

22. Carbone, C.E. et al. Time-resolved cryo-EM visualizes ribosomal translocation with EF-G and GTP. Nature Communications 12, 7236 (2021).

23. Chakraburtty, K. & Kamath, A. Protein synthesis in yeast. International Journal of Biochemistry 20, 581–590 (1988).

24. Ranjan, N. et al. Yeast translation elongation factor eEF3 promotes late stages of tRNA translocation. The EMBO journal 40, e106449 (2021).

25. Qin, S., Xie, A., Bonato, M. & McLaughlin, C. Sequence analysis of the translational elongation factor 3 from Saccharomyces cerevisiae. Journal of Biological Chemistry 265, 1903–1912 (1990).

26. Stark, H., Rodnina, M.V., Wieden, H.-J., Van Heel, M. & Wintermeyer, W. Large-scale movement of elongation factor G and extensive conformational change of the ribosome during translocation. Cell 100, 301–309 (2000).

27. Hassan, A. et al. Ratchet, swivel, tilt and roll: a complete description of subunit rotation in the ribosome. Nucleic Acids Research 51, 919–934 (2023).

28. Flis, J. et al. tRNA translocation by the eukaryotic 80S ribosome and the impact of GTP hydrolysis. Cell reports 25, 2676–2688. e7 (2018).

29. Bourne, H.R., Sanders, D.A. & McCormick, F. The GTPase superfamily: conserved structure and molecular mechanism. nature 349, 117–127 (1991).

30. Taylor, D.J. et al. Structures of modified eEF2· 80S ribosome complexes reveal the role of GTP hydrolysis in translocation. The EMBO journal 26, 2421–2431 (2007).

31. Kaul, G., Pattan, G. & Rafeequi, T. Eukaryotic elongation factor -2 (eEF2): its regulation and peptide chain elongation. Cell biochemistry and function 29, 227–234 (2011).

32. Lucas, B.A. et al. Locating macromolecular assemblies in cells by 2D template matching with cisTEM. Elife 10, e68946 (2021).

33. Wu, C., Huang, X., Cheng, J., Zhu, D. & Zhang, X. High-quality, high-throughput cryo-electron microscopy data collection via beam tilt and astigmatism-free beam-image shift. Journal of structural biology 208, 107396 (2019).

34. You, X. et al. In situ structure of the red algal phycobilisome–PSII–PSI–LHC megacomplex. Nature 616, 199–206 (2023).

35. Zheng, S.Q. et al. MotionCor2: anisotropic correction of beam-induced motion for improved cryo-electron microscopy. Nature methods 14, 331–332 (2017).

36. Rohou, A. & Grigorieff, N. CTFFIND4: Fast and accurate defocus estimation from electron micrographs. Journal of structural biology 192, 216–221 (2015).

37. Wu, S. et al. Diverse roles of assembly factors revealed by structures of late nuclear pre-60S ribosomes. Nature 534, 133–137 (2016).

38. Zivanov, J. et al. New tools for automated high-resolution cryo-EM structure determination in RELION-3. elife 7, e42166 (2018).

39. Punjani, A., Rubinstein, J.L., Fleet, D.J. & Brubaker, M.A. cryoSPARC: algorithms for rapid unsupervised cryo-EM structure determination. Nature methods 14, 290–296 (2017).

40. Pettersen, E.F. et al. UCSF Chimera—a visualization system for exploratory research and analysis. Journal of computational chemistry 25, 1605–1612 (2004).

41. Ben-Shem, A. et al. The structure of the eukaryotic ribosome at 3.0 Å resolution. Science 334, 1524–1529 (2011).

42. Ranjan, N. et al. eEF3 promotes late stages of tRNA translocation on the ribosome. bioRxiv, 20.07. 01.182105 (2020).

43. Bryant, P., Pozzati, G. & Elofsson, A. Improved prediction of protein-protein interactions using AlphaFold2. Nature communications 13, 1265 (2022).

44. Shao, S. et al. Decoding mammalian ribosome-mRNA states by translational GTPase complexes. Cell 167, 1229–1240. e15 (2016).

45. Jørgensen, R. et al. Two crystal structures demonstrate large conformational changes in the eukaryotic ribosomal translocase. Nature Structural & Molecular Biology 10, 379–385 (2003).

46. Svetlov, M.S. et al. Structure of Erm-modified 70S ribosome reveals the mechanism of macrolide resistance. Nature chemical biology 17, 412–420 (2021).

47. Emsley, P. & Cowtan, K. Coot: model-building tools for molecular graphics. Acta crystallographica section D: biological crystallography 60, 2126–2132 (2004).

48. Adams, P.D. et al. PHENIX: a comprehensive Python-based system for macromolecular structure solution. Acta Crystallographica Section D: Biological Crystallography 66, 213–221 (2010).

49. Williams, C.J. et al. MolProbity: More and better reference data for improved all-atom structure validation. Protein Science 27, 293–315 (2018).

