## Supplemental figures for "Snapshots of ribosome dynamics near atomic-resolution in situ: full insight into the eukaryotic elongation cycle"

**Extended Data Figures**

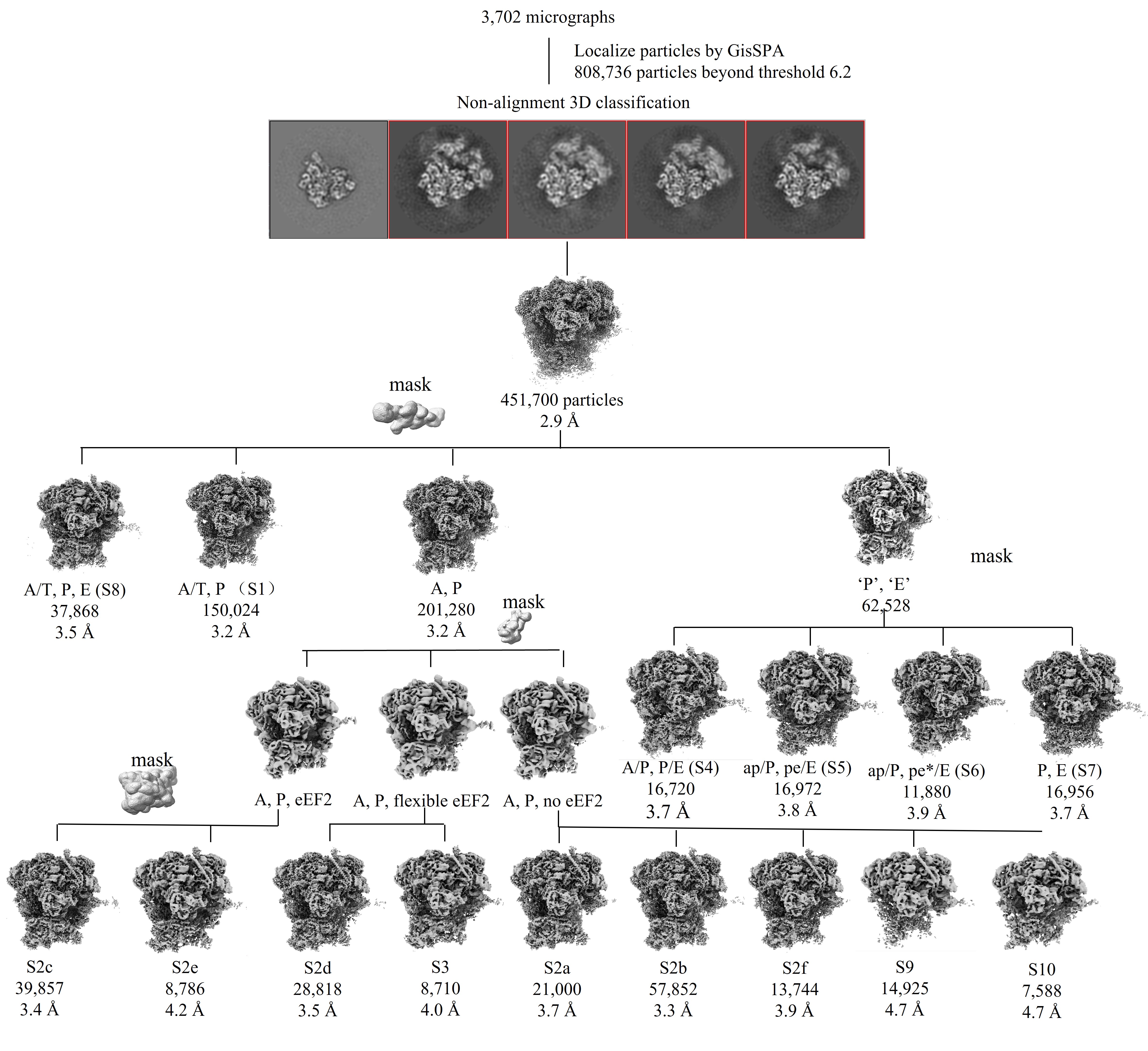

**Extended Data Fig. 1: Image-processing workflow for the identification of the eleven main states.**

The classification process can be divided into four steps: focused classification on the tRNA path region, focused classification on the eEF2 binding region, focused classification on the E-site tRNA binding region, and focused classification on the 40S subunit. The number of particles, corresponding state number, and resolutions are indicated under each structure.

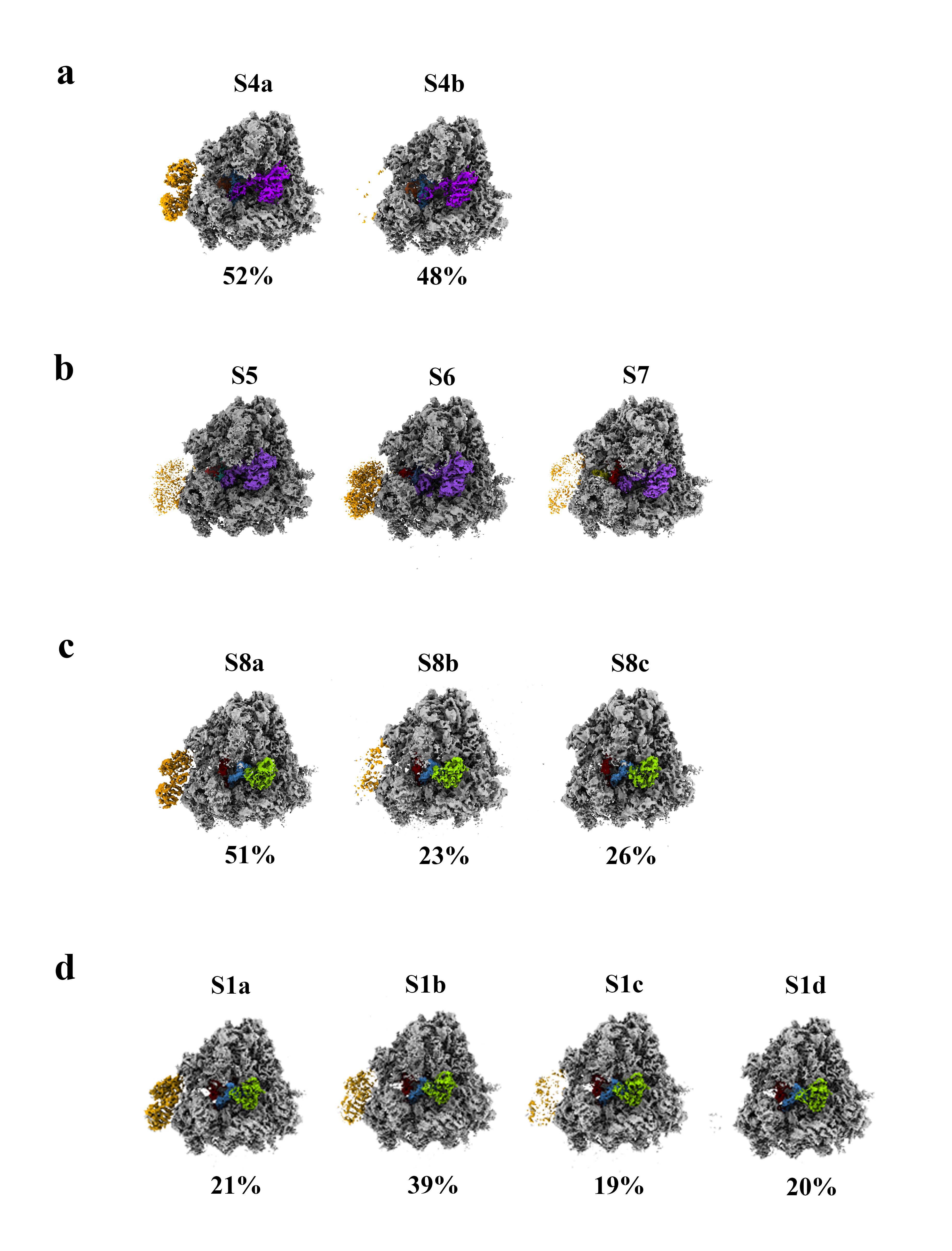

**Extended Data Fig. 2 Exhibitions of the coexistence of eEF3 with eEF2 or eEF1A in different states.**

(**a**) Two sub-states classified from S4 by including eEF3 (S4a, 52%) or not (S4b, 48%). (**b**) The eEF3 shown in S5, S6, and S7. (**c**) Three sub-states divided from ‘A/T, P, E’ state (S8) depending on the presence of eEF3: stable bound to ribosome (S8a), showing flexibility (S8b), and not bound (S8c). The percentages of particles accounted for by each state are listed below the corresponding maps. (**d**) Four sub-states divided from ‘A/T, P’ state (S1) depending on the presence of eEF3: S1a (21%) is the state with stably bound eEF3, S1b (39%) has a little flexibility in eEF3, S1c (19%) shows severer flexibility in eEF3 compared to S1b, and S1d (20%) lost connection with eEF3. The 80S, eEF1A, eEF2 and eEF3 are colored in gray, green, purple and orange, respectively. The tRNAs are colored consistent with Fig. 2.

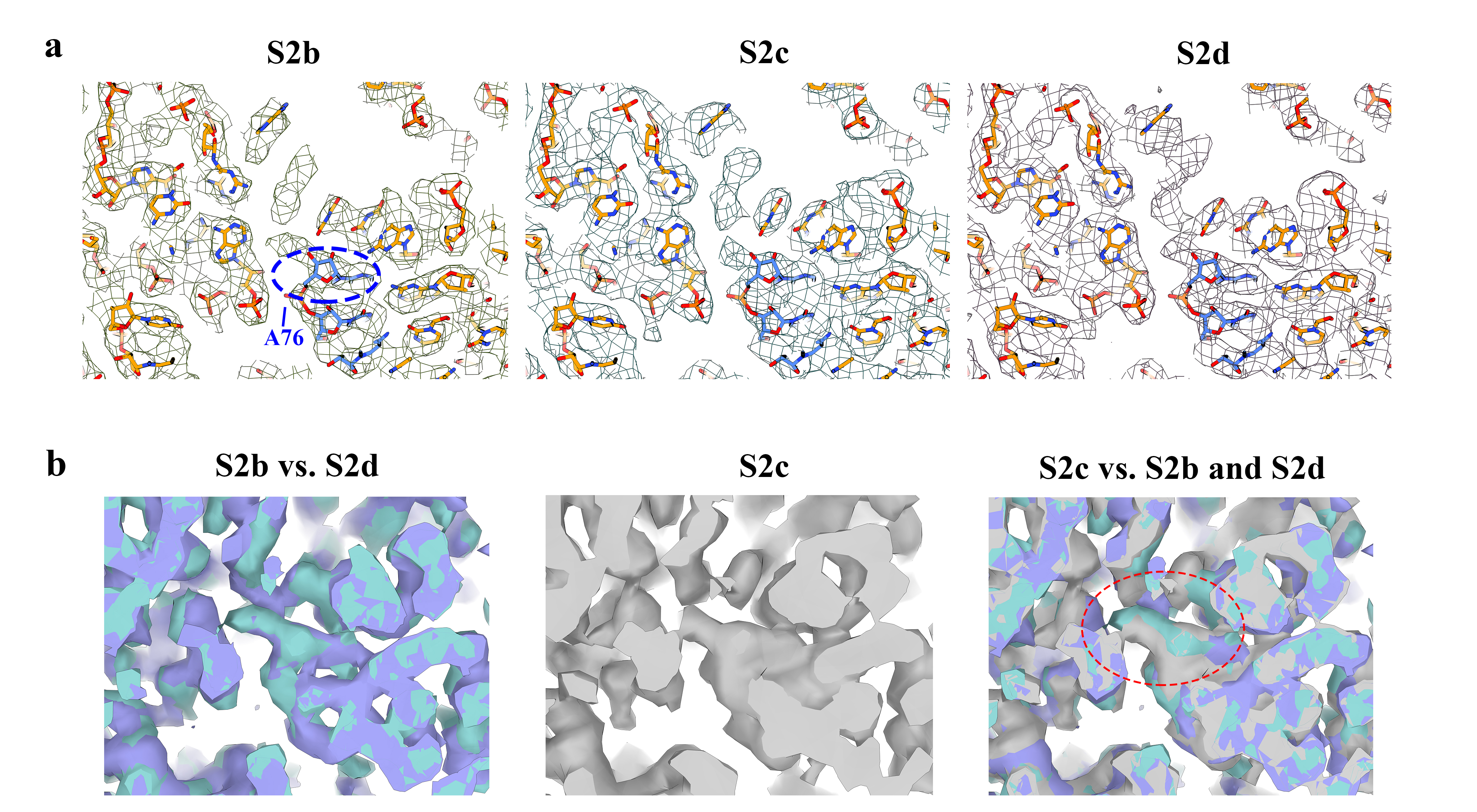

**Extended Data Fig. 3 Density presentation of the three intermediate states (S2b-d) during peptidyl transfer.**

(**a**) The mesh mode presentation depicts the fitting of the atomic models of S2b onto three maps (S2b-d). The atomic models of the A-site tRNA (carbon atoms of model colored in blue) align well with the S2b and S2d maps; however, the fit is less accurate for A76 and the aminoacyl residue in S2c. (**b**) Comparison of the maps among three states. The maps of S2b, S2c and S2d are colored in blue, gray and purple, respectively. The densities of the CCA and aminoacyl residues of the A-site tRNA in S2b and S2d overlap significantly, indicating their similarity. In contrast, the density of the CCA in S2c is less well-resolved compared to the other two states, showing a noticeable shift (highlighted by a red circle) in the superimposed maps.

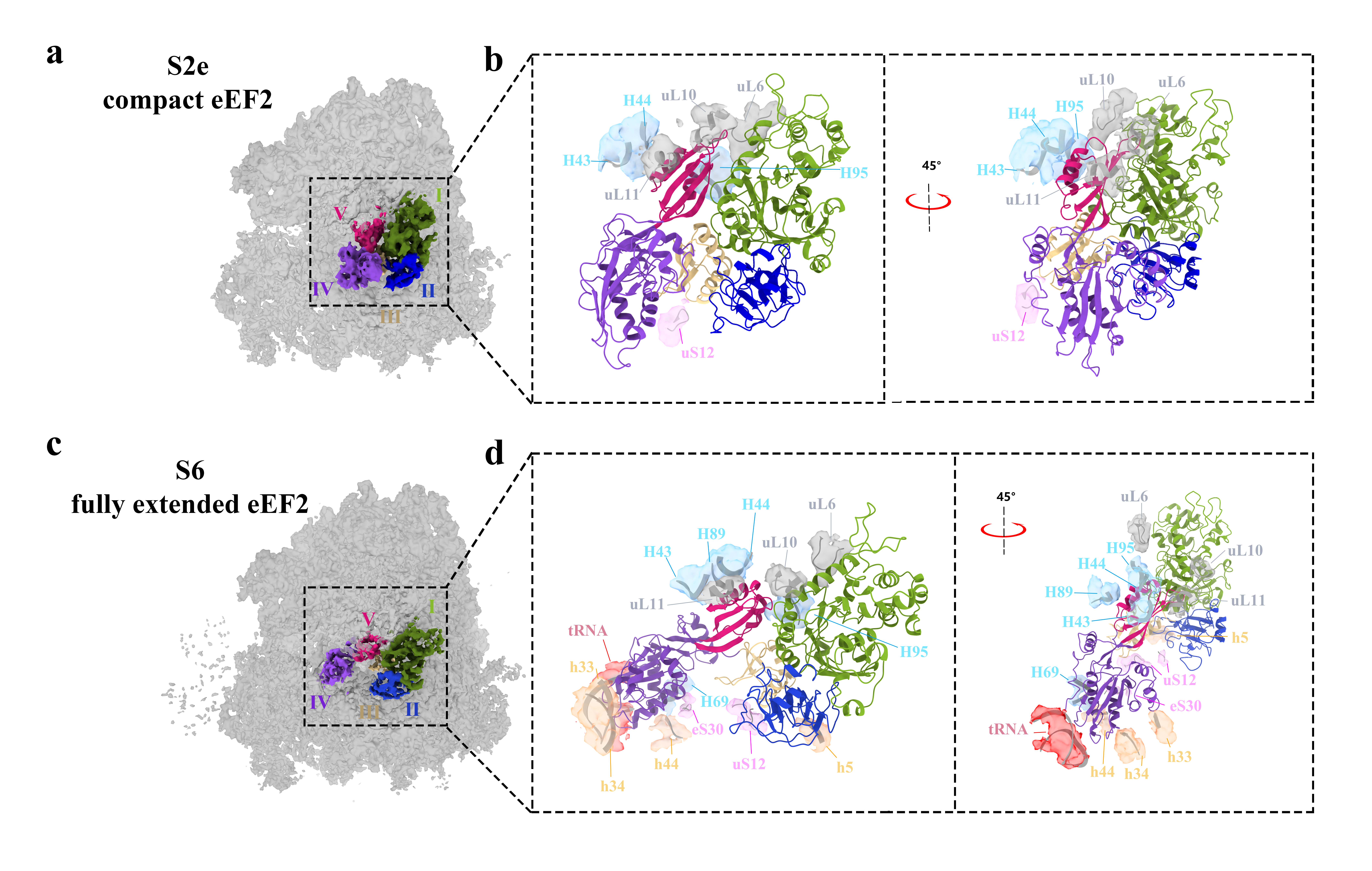

**Extended Data Fig. 4 Surroundings of compact and fully extended eEF2 in S2e and S6.**

(**a**) The overview of compact eEF2 bound to ribosome in S2e. The densities of eEF2 are colored differently: domain I (green), domain II (blue), domain III (yellow), domain IV (purple), and domain V (magenta). (**b**) The environment surrounding compact eEF2 (colored by domain) in the ribosome. Gray and blue densities represent nucleic acids and proteins of LSU, while orange and pink densities represent nucleic acids and proteins of SSU. The models of the ribosome are shown in dark gray ribbon. (**c**) The ­overview of fully extended eEF2 bound to ribosome in S6. (**d**) The environment surrounding of fully extended eEF2 in the ribosome. The arrangement and color are the same as in (b), with the new densities of tRNA interacting with domain IV of eEF2 are colored in red.

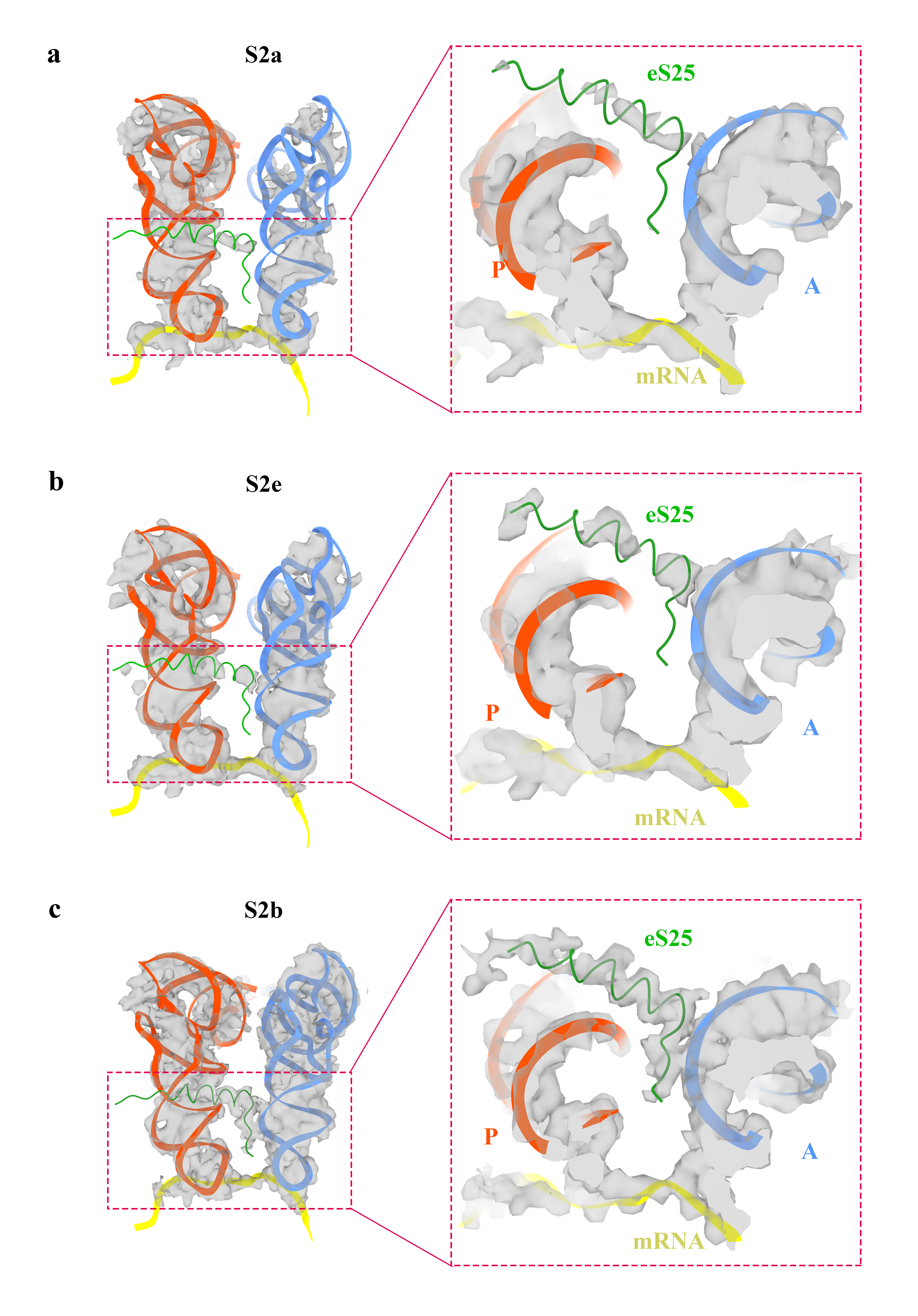

**Extended Data Fig. 5 Density comparison of eS25A helix in different ‘A, P’ states**.

Left panel: Carton diagram depicting A-site tRNA (blue), P-site tRNA (red) and eS25 (green) in S2a (**a**), S2e (**b**) and S2b (**c**). The mRNA is shown in yellow ribbon. Map densities are depicted in gray. Right panel: The close-up view focusing on eS25. The densities of eS25 in S2a (a) and S2e (b) exhibit low quality. In contrast, the helix in S2b (c) shows high continuity with good side chain densities. This helix crosses the P-site tRNA and interacts with the A-site tRNA during peptidyl transfer for comparison.

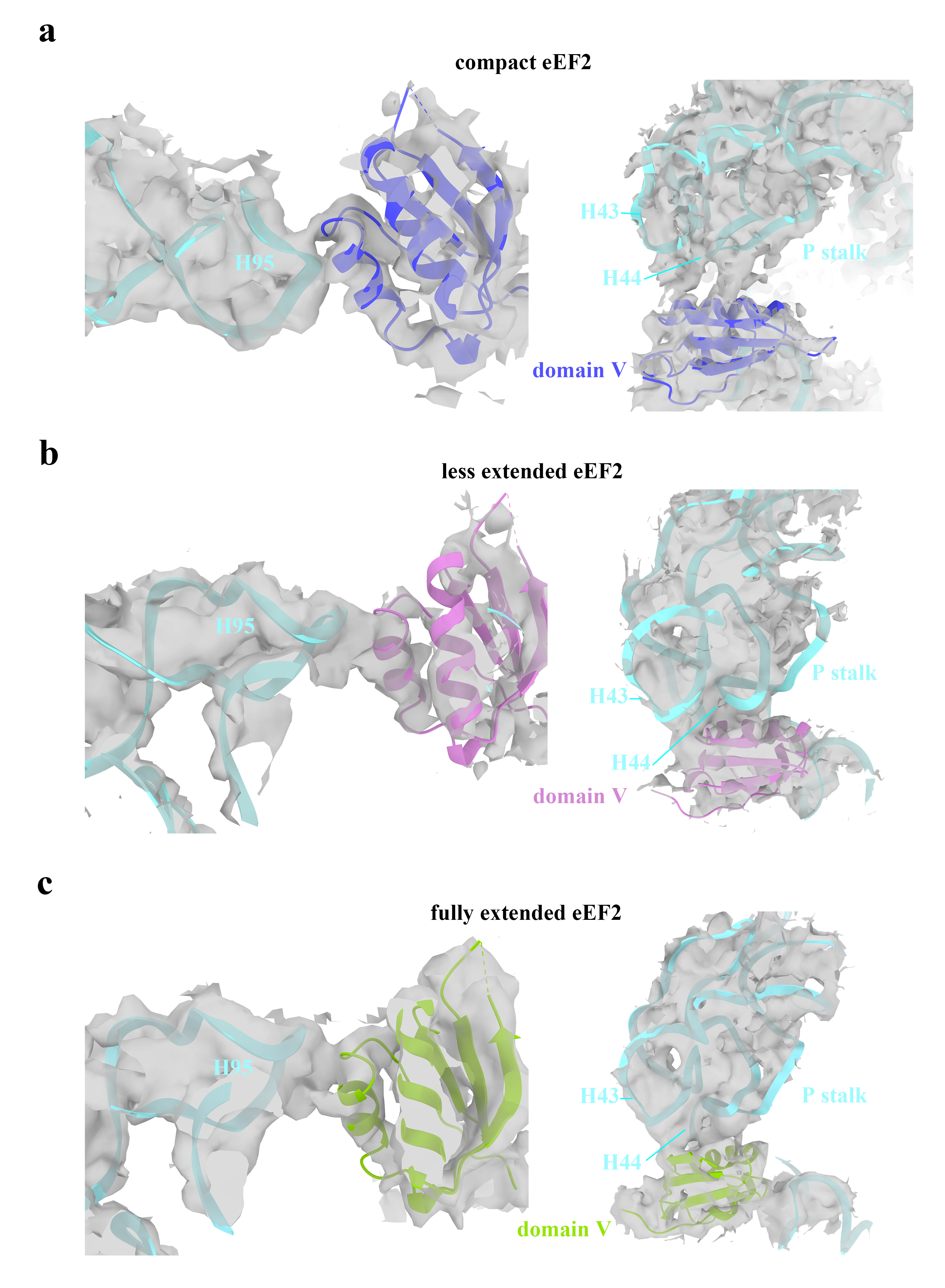

**Extended Data Fig. 6 The interactions of domain V of eEF2 and 25S rRNA in different ribosomal states.**

All maps are represented by gray densities with transparency. The models of H95, H43, and H44 of 25S rRNA are highlighted by cartoon ribbon in blue, while domain V of eEF2 is colored in purple in compact manner (**a**), pink in less-extended manner (**b**), and green in fully extended manner (**c**).

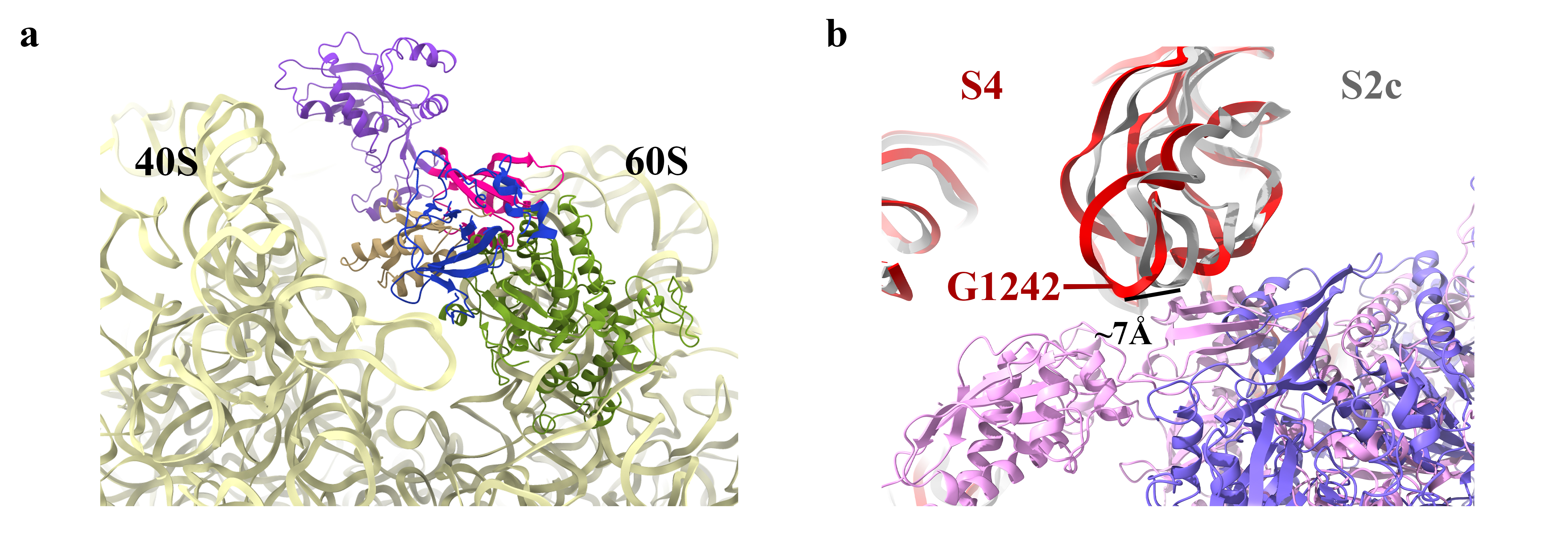

**Extended Data Fig. 7 Superimposing models of compact and extended eEF2 onto the ribosome in S2c.**

(**a**) Incompatibility of fully extended eEF2 with the 80S ribosome (light yellow) of S2c. The models of fully extended eEF2 were positioned by matching domain V (magenta) with compact eEF2 models. Domain I (green) crashes with the 60S subunit because of steric hindrance. Domains II, III, and IV are colored in blue, yellow, and purple, respectively. (**b**) Presentation of P stalk movement of 7 Å from S2c (gray) to S4 (red), with compact eEF2 colored in purple and less extended eEF2 colored in pink.

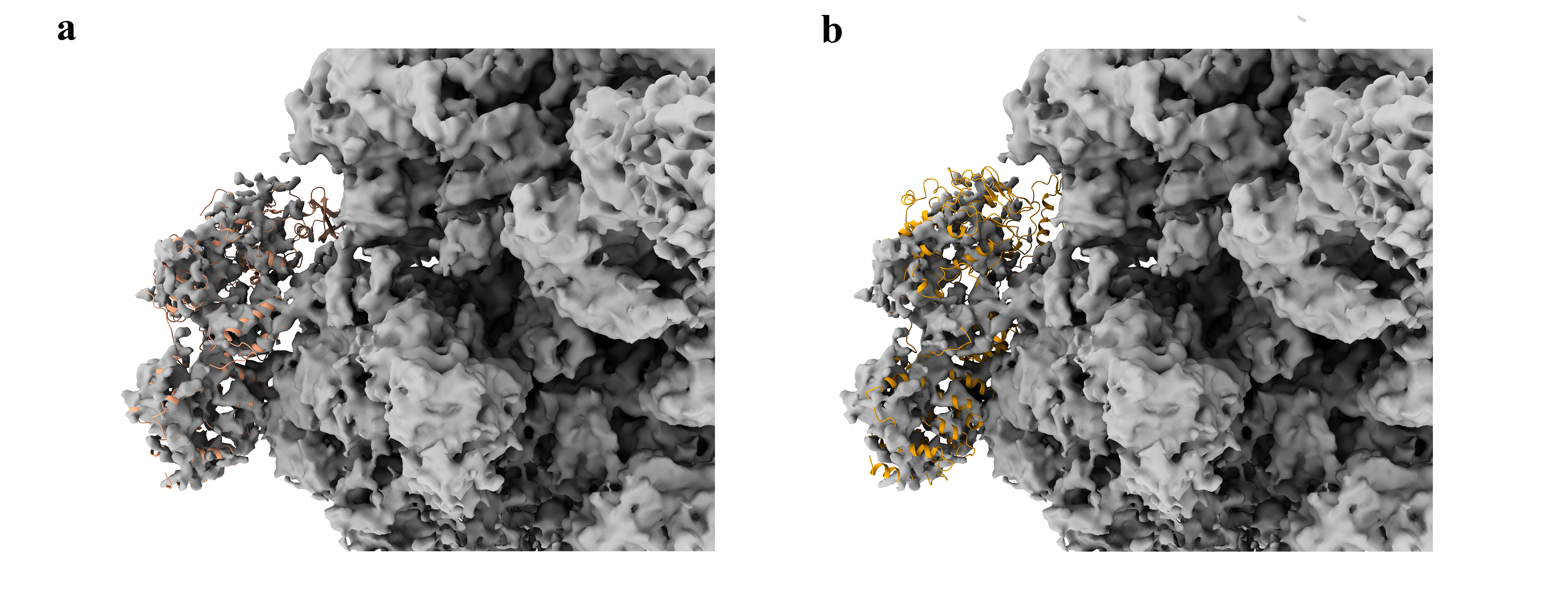

**Extended Data Fig. 8 Atomic models of eEF3 fitted with the corresponding region of EM maps of S6.**

(**a**) The closed-eEF3 models (orange) docking into the EM map (gray) of S6 exhibit a good fit. (**b**) When replacing the models with the open-eEF3, a significant portion of the models extend beyond the boundaries of the EM map. The EM maps were low-pass filtered to 5 Å.

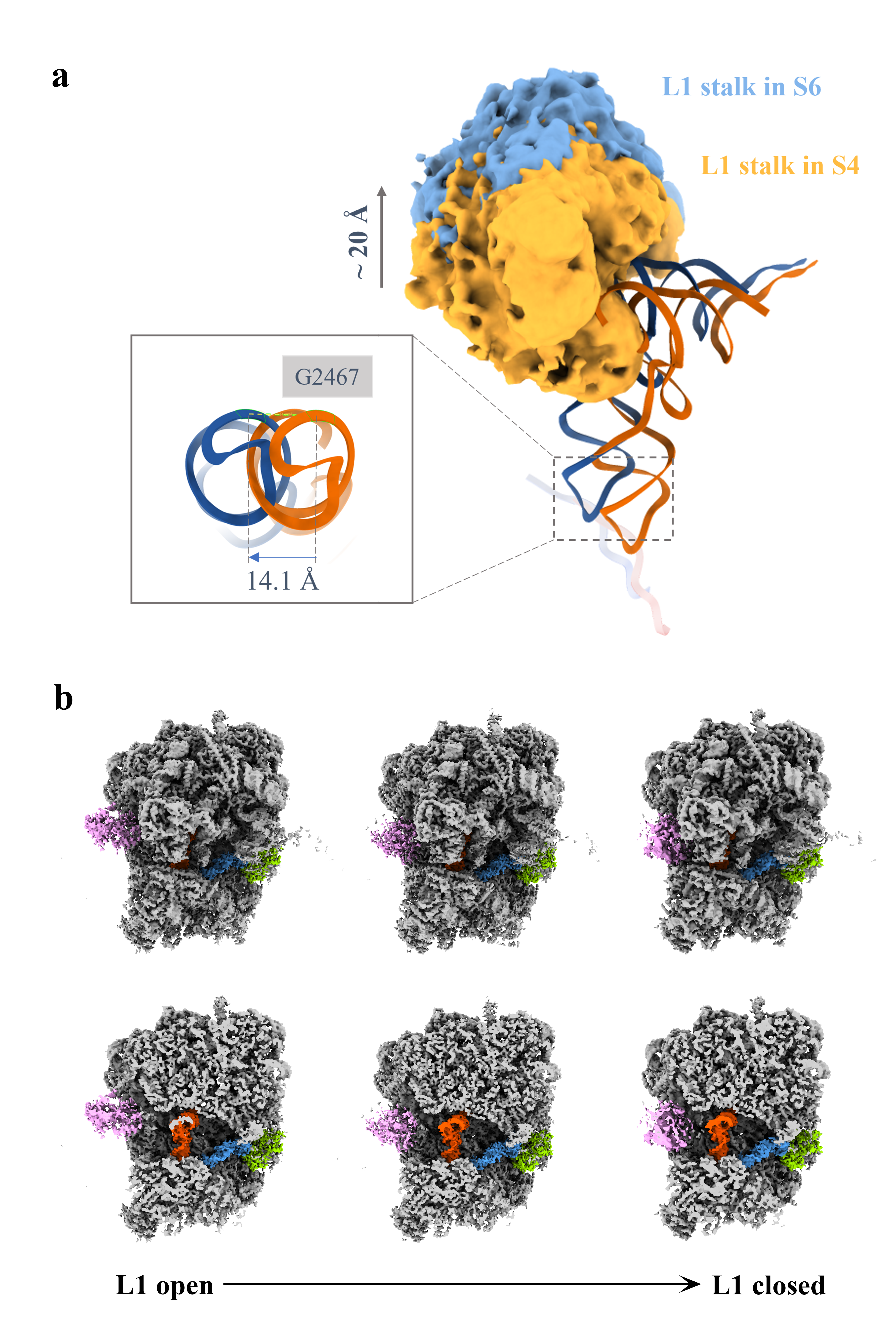

**Extended Data Fig. 9 L1 stalk opens along with the E-site tRNA translocation.**

(**a**) The E-site tRNA undergoes a 14.1 Å movement (left) towards the exit, transitioning from S4 (orange) to S6 (blue), as calculated from the translocation of G2467. Correspondingly, the L1 stalk exhibits an opening of ~20 Å (right), with the L1 stalk densities of S4 and S6 represented by orange and blue, respectively. (**b**) Top row: Transformation of L1 stalk from open to closed in “A/T, P” states obtained by classification on S1 with a focus on the L1 stalk. Bottom row: Corresponding cross-section of the inter-subunit space containing tRNAs. The L1 stalk is colored in pink, and eEF1A is colored in green, A- and P-site tRNAs are highlighted in blue and red, respectively.

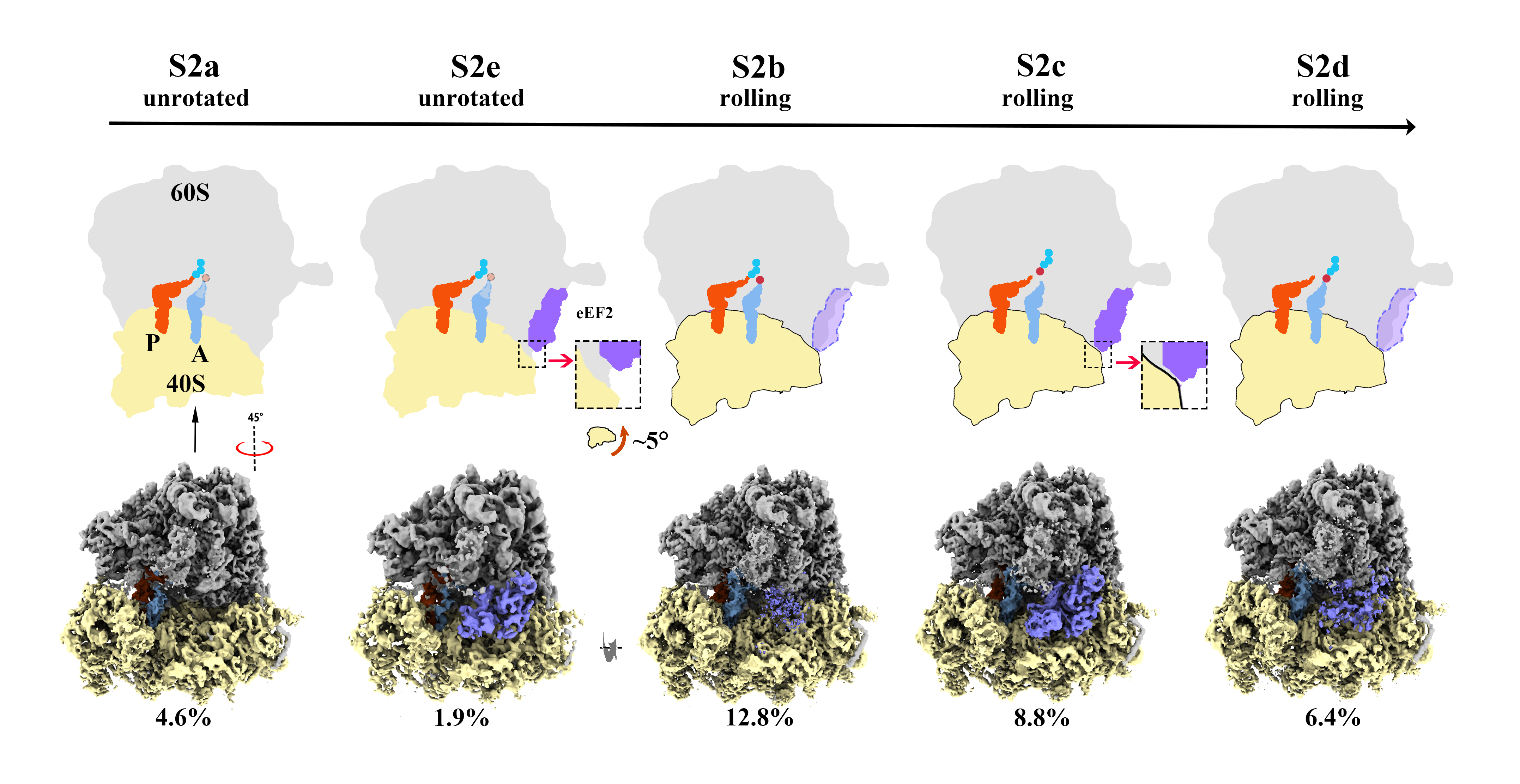

**Extended Data Fig. 10 Hypothesis on kinetic of compact eEF2 during elongation cycle.**

Top row: Cartoons depicting eEF2 conformational changes. The LSU, SSU, and eEF2 are shown in gray, yellow, and purple respectively. The nascent peptidyl chain (blue balls) is attached to the P-site tRNA (red), the aminoacyl residue (red balls) is associated with the A-site tRNA (blue). In the absence of accommodation of the A-site tRNA, the CCA, blurred in the PTC, is outlined by a dotted line and shaded in a lighter color. Stable and flexible binding eEF2 are highlighted in purple with a solid outline and light purple with a dotted outline, respectively. The states of ribosome and movement of SSU are indicated above the corresponding ribosomes. Bottom panels: Corresponding EM map. Percentage of particles accounted for by each state is listed below corresponding ribosomes.
